# Targeting Lysosomal pH Restores Mitochondrial Quality Control in GBA1-Mutant Parkinson’s Disease

**DOI:** 10.1101/2025.08.28.672738

**Authors:** Preethi Sheshadri, Maria Alicia Costa Besada, Alessia Fisher, Szilvia Kiraly, Kritarth Singh, Ioanna Kourouzidou, Thomas S. Blacker, Jialiu Zeng, Orian S Shirihai, Mark W Grinstaff, Michael R Duchen

## Abstract

**Background:** Heterozygous mutations in the Glucocerebrosidase gene (*GBA1*), which encodes the lysosomal enzyme β-glucocerebrosidase (GCase), are a genetic risk factor for Parkinson’s disease (PD). The pathophysiological consequences of *GBA1* mutations on dopaminergic neuronal function, especially their impact on lysosomal function, mitophagy, and mitochondrial bioenergetics, remain unclear.

**Methods:** Fibroblasts and dopaminergic neurons generated from induced pluripotent stem cells (iPSCs) derived from patients with GBA1-PD were used in the study. Live-cell imaging was performed to measure lysosomal acidification, protease activity, mitochondrial membrane potential, and mitophagy. Mitochondrial morphology and autophagic vesicles were examined using transmission electron microscopy. Oxygen consumption rate was measured by Seahorse assay. V-ATPase assembly was quantified using FLIM-FRET, and pharmacological interventions included rapamycin and acidic nanoparticles. Statistical analyses involved unpaired t-tests, one-way ANOVA, and two-way ANOVA.

**Results:** GCase activity, lysosomal acidification, protease activity, mitophagy and mitochondrial bioenergetic function were all impaired. Mitochondria were fragmented, with reduced membrane potential and oxygen consumption. MTORC1 was constitutively phosphorylated and FLIM-FRET measurements confirmed impaired lysosomal V-ATPase assembly, which was reversed following rapamycin treatment. Rapamycin and lysosome-targeted acidic nanoparticles rescued lysosomal pH, restored mitophagy, mitochondrial membrane potential and mitochondrial OXPHOS complex levels in *GBA1* mutant dopaminergic neurons.

**Conclusions:** We reveal a novel mechanistic link between *GBA1* mutations and mitochondrial dysfunction, as disruption of V-ATPase assembly driven by MTORC1 activation impairs lysosomal acidification. Mitophagy is therefore impaired leading to mitochondrial dysfunction, undermining dopaminergic cell function and fate. Pharmacological intervention with rapamycin or acidic nanoparticles restore lysosomal pH and rescue mitochondrial function, signposting a novel therapeutic approach for GBA1-PD.

## Introduction

Genome-wide association studies have revealed that mutations of the glucocerebrosidase (*GBA1*) gene constitute a significant risk factor in the development of Parkinson’s Disease (PD)[1]. The *GBA1* gene encodes the enzyme β-glucocerebrosidase (GCase), which generates glucose and ceramide from glucosylceramide within lysosomes. While homozygous *GBA1* mutations cause Gaucher Disease, which may include a significant neurodegenerative component[2], heterozygous mutations are associated with an increased risk of developing PD. The two most prevalent *GBA1* mutations associated with PD are the N370S and L444P mutations[3]. Intriguingly, the less common E326K mutation in *GBA1* exhibits a relatively mild effect on GCase activity, does not cause Gaucher’s Disease, but is correlated with PD risk[4].

Mitochondrial dysfunction appears to constitute a defining characteristic of PD[5]. The adverse impact of dysfunctional mitochondrial oxidative phosphorylation (OXPHOS) on the viability of dopaminergic (DA) neurons is substantiated by experimental models of PD that result from toxins that target complex I[5]. The *Gba1* mouse knockout model, a model of severe neurodegeneration, exhibited severe mitochondrial dysfunction in primary neurons and astrocytes in culture, alongside neurological pathologies associated with PD, including disruption of autophagy-lysosomal pathways, and the accumulation of ubiquitinated proteins and α-synuclein[6].

The autophagosome-lysosome axis and ubiquitin proteasome systems accomplish degradation of dysfunctional organelles and protein degradation and together play a critical role in cellular quality control[7]. Lysosomal acidification generating pH values between 4.5 to 4.7[8] is essential for normal lysosomal function and is required for the activity of lysosomal hydrolytic enzymes and effective protein degradation[44,12]. The acidification is generated by the vacuolar-type H^+^ ATPase (V-ATPase), a proton pump composed of a peripheral V_1_ domain, which hydrolyses ATP, and a membrane-integrated V_o_ domain, responsible for the translocation of protons into the lysosomal lumen[10]. Disrupted V-ATPase assembly and V-ATPase dysfunction have been reported associated with multiple disorders[11], including Juvenile-onset Parkinson’s Disease[12], Alzheimer’s Disease[13,14], Epilepsy[15,16] and Down’s syndrome[17].

*GBA1*-linked pathologies such as Gaucher’s disease and PD are associated with a dysfunctional autophagy-lysosome pathway, including impaired lysosomal regeneration from autolysosomes during macroautophagy[18,19]. These investigations suggest that targeting the dysfunctional lysosomal system and the autophagy-lysosome pathway may serve as a potential therapeutic strategy for PD linked to *GBA1* mutations. Although numerous studies indicate that impaired lysosomal acidification plays a role in neuronal pathologies associated with various neurodegenerative disorders[38] and that GBA1-PD is characterised by lysosomal dysfunction [44], the effects of compromised lysosomal acidification in GBA1-PD have not been extensively examined.

In this study, we have characterised the consequences of *GBA1* mutations associated with PD for mitochondrial and lysosomal function in patient-derived fibroblasts and DA neurons generated from patient-derived induced pluripotent stem cells (iPSCs). Since GCase is a lysosomal enzyme, we hypothesised that lysosomal dysfunction caused by the mutations might impair mitochondrial function, initiating a pathophysiological cascade culminating in cellular malfunction. We also discovered that constitutive and inappropriate activation of mechanistic target of rapamycin complex 1 (MTORC1) impairs lysosomal acidification by disrupted assembly of the lysosomal V-ATPase complex, resulting in lysosomal dysfunction. Additionally, we demonstrate that restoring lysosomal pH by treatment with rapamycin or acidic nanoparticles (acidic NPs) targeted to lysosomes enhances mitophagy and rescues both lysosomal and mitochondrial function in cells carrying GBA1-PD mutations.

## Results

To characterise the impact of PD-related *GBA1* mutations on lysosomal and mitochondrial function, we examined three lines of human dermal fibroblasts and six iPSC lines derived from GBA1-PD patients carrying E326K or N370S mutations, along with two healthy controls. We also included one CRISPR-corrected isogenic control iPSC line each for GBA1-N370S and GBA1-E326K iPSCs. (Table 1). We first performed the experiments in fibroblasts as a proof of principle, and subsequently confirmed the key findings in iPSC-derived DA neurons.

**Table 1.** List of cell lines used.

| Cell Lines used | Cell Line ID | Age/Sex | Cell type | Mutation | Source | Ethics Committee Approval |
| --- | --- | --- | --- | --- | --- | --- |
| CONTROL 1 | CONTROL 1 | 77/Male | Fibroblasts | No Mutation | NHNN | PD Royal Free Ethics |
| N370S | ND34263 | 65/Male | Fibroblasts; source for NH50182 iPSCs | GBA1-N370S; Heterozygous | NIH-RUCDR |  |
| CONTROL 2 | CONTROL 2 | 55/Male | Fibroblasts; source for Control1 iPSCs | No Mutation | NHNN | PD Royal Free Ethics |
| E326K 1 | Het1/ JS48753 | 58/Male | Fibroblasts; source for Het1/E326K 1 iPSCs | GBA1-E326K; Heterozygous | NHNN | PD Royal Free Ethics |
| E326K 2 | ND41015 | 63/Male | Fibroblasts; source for ND50045 iPSCs | GBA1-E326K; Heterozygous | NIH-RUCDR |  |
| Isogenic Control (IsoC) | NH50142 | 63/Male | iPSCs; Isogenic Control for ND50045 | CRISPR corrected GBA1-E326K heterozygous mutation | NIH-RUCDR |  |
| E326K | ND50045 | 63/Male | iPSCs | GBA1-E326K; Heterozygous | NIH-RUCDR |  |
| Isogenic Control (IsoC) | NH50186 | 65/Male | iPSCs; Isogenic Control for NH50182 | CRISPR corrected GBA1-N370S heterozygous mutation | NIH-RUCDR |  |
| N370S | NH50182 | 65/Male | iPSCs | GBA1-N370S; Heterozygous | NIH-RUCDR |  |
| Control | Control 1 | 55/Male | iPSCs | No Mutation | Derived from CONTROL 2 fibroblasts | PD Royal Free Ethics for CONTROL 2 fibroblasts |
| E326K 1 | Het1/ JS48753 | 58/Male | iPSCs | GBA1-E326K; Heterozygous | Derived from Het1 fibroblasts | PD Royal Free Ethics for JS48753 fibroblasts |
| E326K 2 | Het 2/ MB240649 | 63/Male | iPSCs | GBA1-E326K; Heterozygous | Derived from MB240649 fibroblasts | PD Royal Free Ethics (for MB240649 fibroblasts) |
| Control | SFC156 (Control 2) | 65/Male | iPSCs | No Mutation | EBiSC |  |
| N370S 1 | SFC834 (N370S1) | 72/Male | iPSCs | GBA1-N370S; Heterozygous | EBiSC |  |
| N370S 2 | SFC848 (N370S2) | 68/Male | iPSCs | GBA1-N370S; Heterozygous | EBiSC |  |

### Lysosomal function is impaired in cells with PD-related *GBA1* mutations

As GCase is a lysosomal enzyme, our investigation began by characterising the lysosomes in fibroblasts and DA neurons from both control and PD patient-derived iPSCs. The GBA1 protein expression levels measured by western blot were not significantly different between the control and *GBA1* mutant neurons; however, GCase enzymatic activity was significantly reduced in iPSC-DA neurons and fibroblasts carrying either *GBA1* mutation (Fig 1A, B, S1A, B). Lysosomal acidification was measured using the ratiometric pH-sensitive dye, Lysosensor Yellow/Blue DND160. These data revealed impaired acidification in both E326K and N370S DA neurons and fibroblasts (Fig 1C, S1C, D). Calibration of the lysosomal pH (see methods) gave values of 4.74±0.03 - 4.77±0.02 in control cells but between 4.89±0.02 and 5.06±0.03 in the mutant fibroblasts (Fig 1D). The function of endocytic trafficking and lysosomal proteolytic activity was measured using the DQ-Red BSA assay. DQ-Red BSA intensity was significantly reduced in the *GBA1* mutant neurons, indicating impaired lysosomal degradative function in GBA1-PD cells (Fig 1E, S1E). Autofluorescence excited at 355nm and imaged between 400-600nm showed an unusually bright extramitochondrial component in GBA1-PD fibroblasts (Fig 1F) that was significantly greater than control cells. Spectral scanning and linear unmixing separated the expected mitochondrial NADH signal with an emission peak at 450nm and an extensive non-mitochondrial signal with a peak emission at 480nm, attributed to accumulated lipofuscin. Lipofuscins are cytoplasmic granules generated as a consequence of autophagy and phagocytosis processes, which show auto-fluorescence over a broad spectrum ranging from 480nm to 700nm when excited by ultraviolet or blue light[20]. While the accumulation of lipofuscin serves as an indicator of ageing, abnormal accumulation is consistent with impaired lysosomal acidification and defective autophagy[21]. Accumulation of lipofuscin is also a hallmark of Batten Disease (neuronal ceroid-lipofuscinoses), a severe early-onset neurodevelopmental disorder with progressive neurodegeneration also associated with lysosomal dysfunction [22]. These data collectively indicate significant lysosomal dysfunction in the GBA1-PD cells.

**Figure 1:**
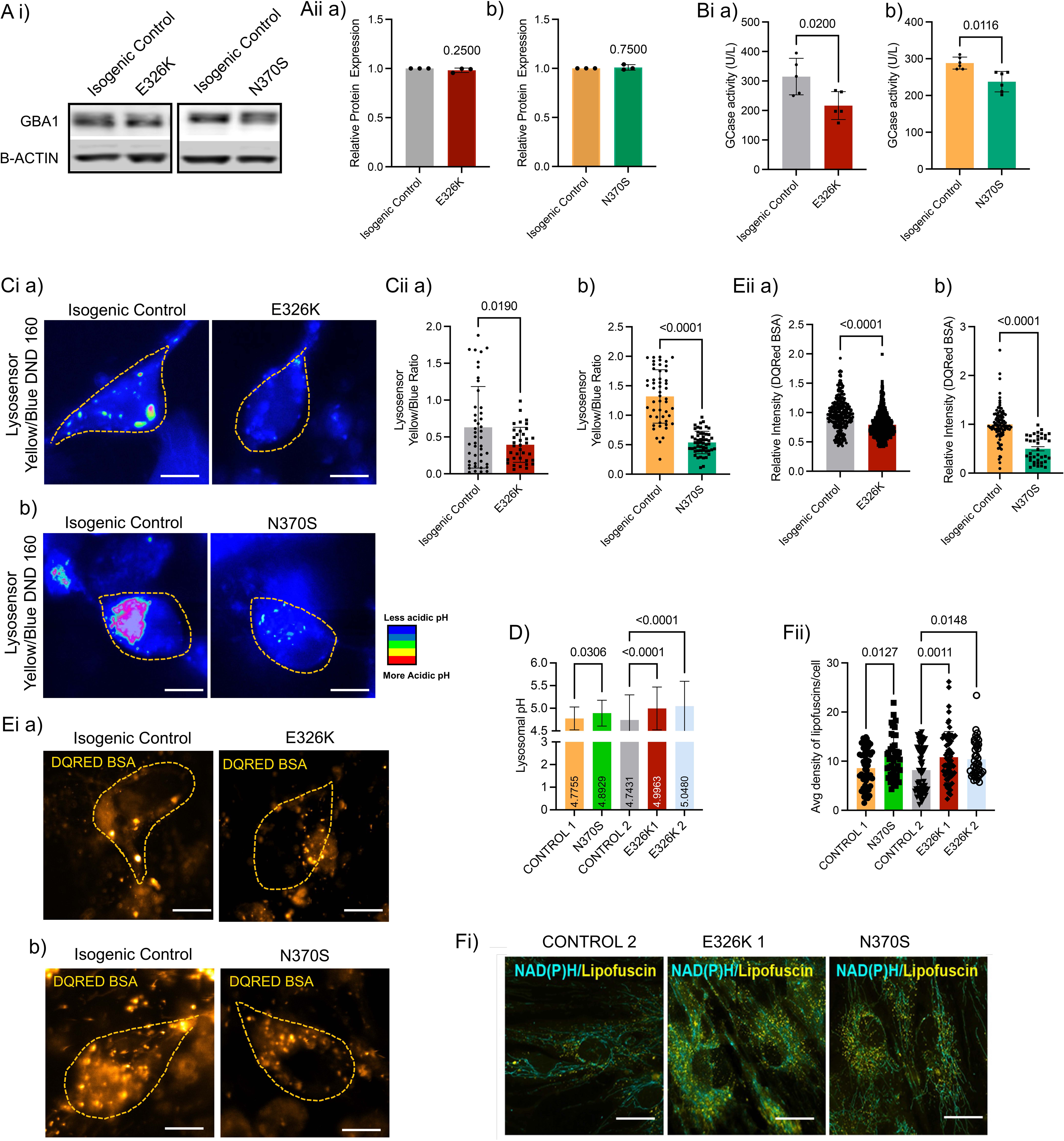
Lysosomal abnormalities in GBA1-PD DA cells. Representative western blot images of GBA1 protein expression (Ai) and quantification (Aii a,b) in isogenic control and GBA1-E326K and N370S DA neurons. GCase activity in GBA1-E326K (Bi a) and GBA1-N370S (Bi b) DA neurons and respective isogenic controls. Representative ratioed images (Ci a, b) and quantification of fluorescence ratio of GBA1-E326K (Cii a) and GBA1-N370S (Cii b) DA neurons and respective isogenic controls labelled with Lysosensor Yellow/Blue DND 160, indicating lysosomal pH in cells. D) Histogram depicting mean lysosomal pH values in *GBA1* mutant fibroblasts. Representative confocal images (Ei a,b) and quantification of relative intensity of GBA1-E326K (Eii a) and GBA1-N370S (Eii b) DA neurons and respective isogenic controls stained with DQ-Red BSA indicative of proteolytic activity. Representative confocal images (Fi) and quantification (Fii) of lipofuscins in GBA1 fibroblasts, which are seen upon autofluorescence. Data represented as mean±SD; one-sample Wilcoxon rank t-test, unpaired t-test or One-way ANOVA with Kruskal-Wallis and Dunn’s multiple comparisons test. p-values are noted on the graphs; the statistical tests, corresponding effect sizes, and confidence intervals for each graph are in supplementary table 2.

### Mitochondria are dysfunctional in cells with GBA1-PD mutations

To determine whether the impaired lysosomal activity in the *GBA1* mutations has a secondary impact on mitochondrial form and function, we measured mitochondrial membrane potential (ΔΨm) using the potentiometric fluorescent reporter tetramethyl-rhodamine methyl ester (TMRM). ΔΨm was significantly reduced in E326K and N370S *GBA1* mutant DA neurons and fibroblasts (Fig 2A, S2A, C). The lower ΔΨm was accompanied by mitochondrial fragmentation in the N370S and E326K DA neurons and fibroblasts (Fig 2Aii c,d, S2Aii b). Ultrastructure analysis using electron microscopy revealed significant differences in mitochondrial morphology in the *GBA1* mutant neurons, including reduced cristae density and increased mitochondrial area, along with decreased mitochondrial aspect ratio, indicating mitochondrial swelling and fragmentation in the *GBA1* mutant iPSC DA neurons (Fig. 2B). Measurements of oxygen consumption rates using the ‘Seahorse’ respirometry system revealed ∼40% decrease in basal oxygen consumption rate in the *GBA1* mutant DA neurons with the isogenic controls (Fig 2C). To exclude changes in mitochondrial mass and to explore the substrate dependence of mitochondrial respiration, we used mitochondria isolated from the cultures. This allowed measurements of respiratory rates with different substrates, favouring complex I (pyruvate/malate) or complex II (rotenone/succinate). ADP-stimulated respiration (State 3) rate using both CI and CII-dependent substrates was significantly reduced in the *GBA1* mutant neurons compared to the control neurons (Fig 2D, S2C). Western Blots of the OXPHOS complex proteins using a cocktail of antibodies to respiratory chain proteins, revealed a significant increase in the expression of complex IV in E326K DA neurons (Fig 2Eii a and S2Dii a) and increased expression of complex I proteins in N370S neurons compared to the isogenic controls (Fig 2Eii b and S2Dii b). Complex III was reduced in both E326K and N370S DA neurons compared to the isogenic controls while the expression of other respiratory chain proteins remained unaltered. None of these data suggested any defect in complex I assembly or function. These data collectively point to impaired mitochondrial bioenergetic function in fibroblasts and neurons carrying *GBA1* mutations.

**Figure 2:**
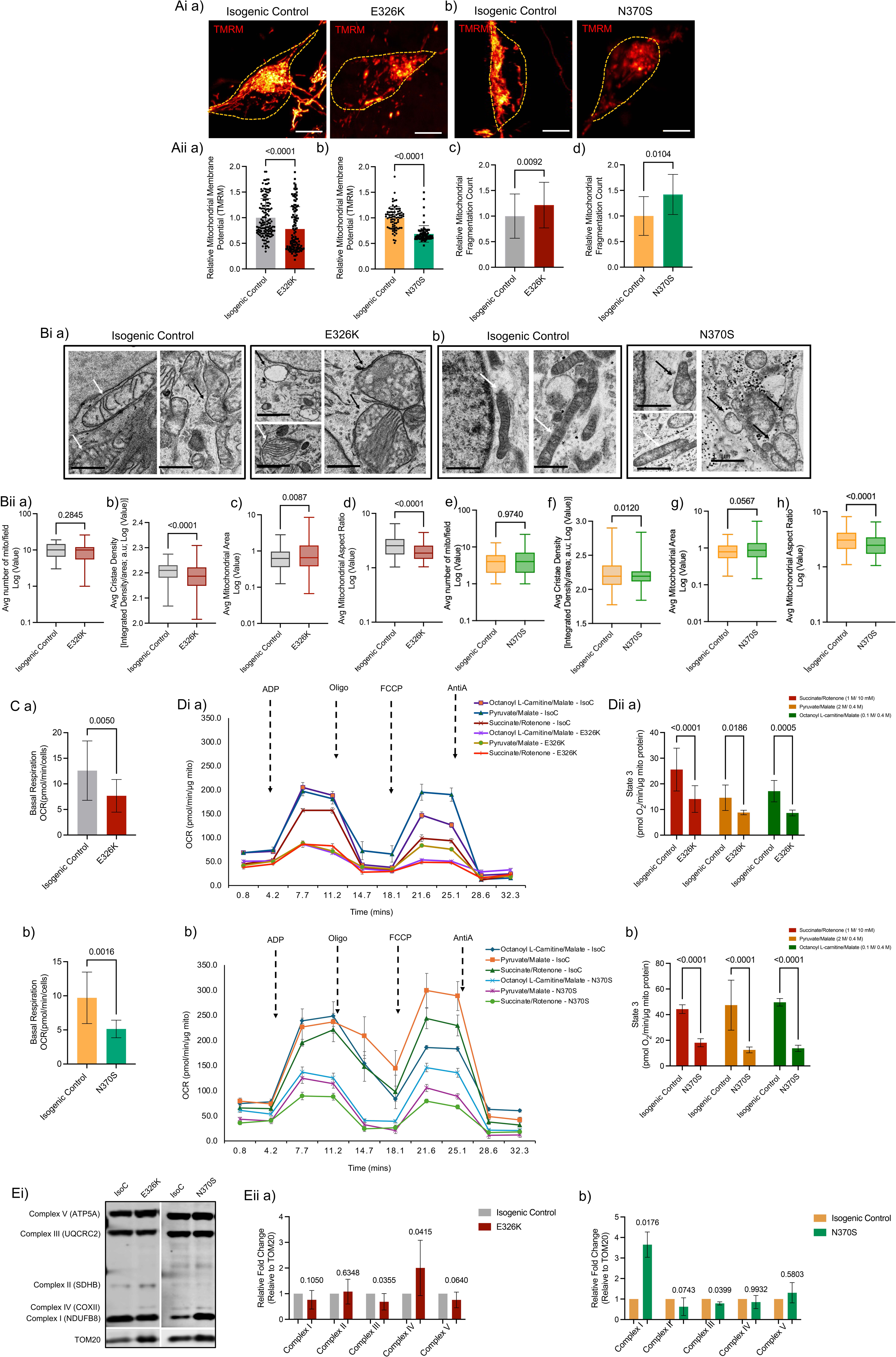
Mitochondrial dysfunction in GBA1-PD DA neurons. Representative confocal images of GBA1-E326K (Ai a) and GBA1-N370S (Ai b) DA neurons along with isogenic controls stained with TMRM and subsequent quantification of fluorescent intensity to measure ΔΨm (Aii a, b) and mitochondrial fragmentation count (Aii c,d). Representative ultramicroscopic images of isogenic control and GBA1-E326K (Bi a) and GBA1-N370S (Bi b) DA neurons. Bii– Box-and-whisker plots showing the minimum, maximum, median, and interquartile range of the log(Y) values of the morphological parameters - average number of mitochondria(a, e), average cristae density in mitochondria (b, f), average mitochondrial area (c, g) and mitochondrial aspect ratio (d, h). The white arrow indicates normal mitochondria; the black arrow indicates abnormal mitochondria. Histograms representing the basal respiration rate in *GBA1* mutant iPSC-DA neurons from GBA1-E326K (C a) and GBA1-N370S mutant lines(C b) and respective isogenic controls. Representative seahorse plots showing oxygen consumption rates in iPSC-DA neurons from GBA1-E326K (Di a) and GBA1-N370S (Di b) DA neurons and respective isogenic controls, using various substrates. Histograms representing State 3(ATP Production) in mitochondria isolated from GBA1-E326K (Dii a) and GBA1-N370S (Dii b) DA neurons and isogenic controls, as measured by seahorse assay. E i and ii) Representative immunoblotting images (Ei) with subsequent quantification of OXPHOS complex proteins in GBA1 E326K (Eii a) and N370S (Eii b) DA neurons. Data represented as mean±SD; Statistics: Mann-Whitney’s test, log-transformed and unpaired t-test, one-sample Wilcoxon rank test or Ordinary two-way ANOVA with Šidák multiple comparisons test with single pooled variance. p-values are noted on the graphs. The statistical tests, corresponding effect sizes, and confidence intervals for each graph are reported in supplementary table 2.

### Mitophagy is impaired in cells with GBA1-PD mutations

We then set out to explore the underlying mechanisms that link impaired lysosomal function to impaired mitochondrial bioenergetics in the GBA1-PD fibroblasts and DA neurons. A logical mechanism linking impaired lysosomal function with mitochondrial dysfunction might operate through dysfunctional mitophagy and our earlier work also established dysfunctional mitophagy in *Gba1* KO mice[6]. We quantified mitophagy using the dual excitation probe mt-Keima (Fig 3A). mt-Keima is a pH-sensitive probe that measures the fraction of mitochondria in neutral (pH 7 - 7.8 in the mitochondrial matrix) versus acidic pH (pH 4.5-4.7 in lysosomes) environment[23]. The mt-Keima signal ratio was significantly reduced in fibroblasts and DA neurons in the fibroblasts and DA neurons carrying both the GBA1-N370S and GBA1-E326K mutations indicating impaired mitophagy (Fig 3Aii. S3Aii, Bii). Western blotting to quantify the status of autophagy pathways in the *GBA1* mutant neurons revealed increased LAMP1 levels in E326K and N370S neurons, along with increased pMTORC1/MTORC1 ratio in the two mutant types. While LC3 flux, and expression of p62 and TOM20 were increased in the E326K mutant iPSC DA neurons, these were not significantly altered in the N370S DA neurons (Fig 3B). Since the lysosomal pH is increased in GBA1-PD cells (Fig 1D), the reduced mt-Keima signal in GBA1-PD cells could be attributed either to the altered lysosomal pH or to the impaired fusion of autophagosomes to lysosomes. To address this, we performed triple staining of fibroblasts against citrate synthase (CiS, for mitochondria), LAMP1 (for lysosomes), and LC3 (for autophagosomes). Subsequent imaging revealed that although there was no significant difference in mitophagosome (mitochondria and autophagosome colocalisation) density between *GBA1* mutant and control fibroblasts, mitolysosome (colocalisation of mitochondria and lysosomes) density was significantly increased in the *GBA1* mutant fibroblasts (Fig 3C). Characterisation of electron micrographs for specific autophagic vesicle types based on Neikirk et al.,2023[24], showed that autophagic vesicles, specifically the autolysosome and lysosome number, were significantly increased in both *GBA1* mutant fibroblasts and neurons. (Fig 3Dii, S3Cii a and b). From the micrographs, the autophagosomes also appeared to be fully enclosed, ruling out incomplete phagophore formation as the underlying mechanism of autophagy defect. These data indicated that mitophagy is significantly impaired in cells carrying the GBA1-PD mutations and the decrease in mitophagy is likely due to impaired pH and not improper phagophore formation.

**Figure 3:**
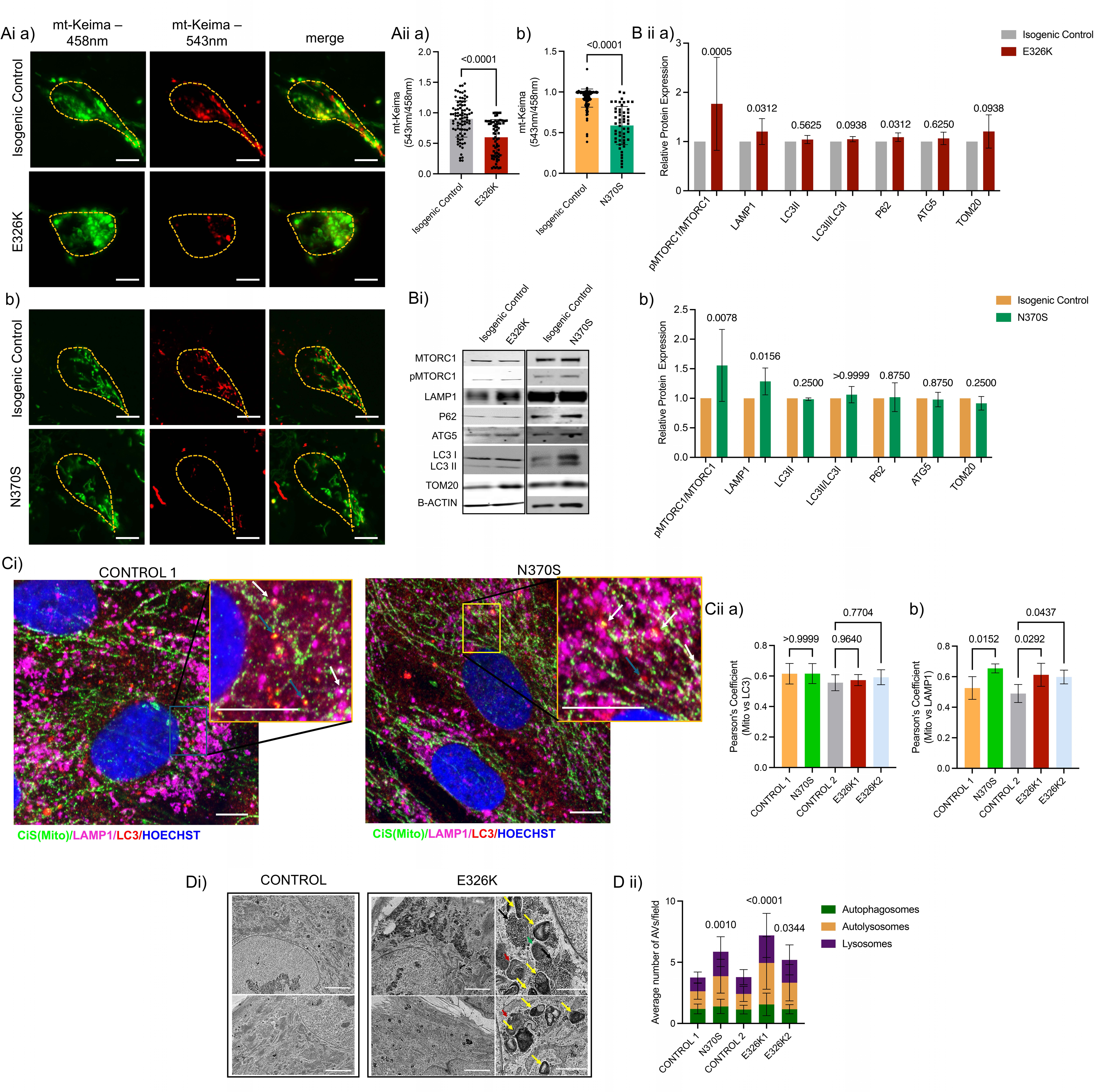
Mitophagy defects in GBA1-PD DA neurons. Representative images of GBA1-E326K (Ai a)and GBA1-N370S (Ai b) DA neurons and isogenic controls transduced with mt-Keima plasmid and imaged with excitation at 458 and 543nm and subsequent quantification of the ratio of signals at 543/458nm (Aii a,b). Representative western blot images of *GBA1* mutant and isogenic control lysates probed for autophagic proteins (Bi) and subsequent quantification of protein levels (Bii a,b) Representative Immunofluorescence images of control and N370S fibroblasts probed against Citrate Synthase (CiS), LAMP1 and LC3, labelling mitochondria, lysosomes and autophagosomes respectively (Ci) and subsequent quantification of colocalisation co-efficient of mitophagosomes (Cii a) and mitolysosomes(Cii b). Representative TEM images (Di) and quantification (Dii) of control and E326K mutant fibroblasts capture different autophagy phases in the cells. Black arrows may indicate protein aggregates, yellow arrows indicate autolysosomes, red arrows indicate autophagosomes, and green arrows indicate lysosomes in the cells. Data represented as mean±SD; Statistics: Mann-Whitney’s test, one-sample Wilcoxon rank test, One-way ANOVA with Holm-Šidák multiple comparisons test or Ordinary two-way ANOVA with Šidák multiple comparisons test. p-values are noted on the graphs. The statistical tests, corresponding effect sizes, and confidence intervals for each graph are in supplementary table 2.

### V-ATPase complex formation is impaired in GBA1-PD

Since our data showed an accumulation of lysosomes and autolysosomes in GBA1-PD fibroblasts and iPSC-DA neurons, we suspected this was a result of impaired lysosomal acidification. Assembly of the pH regulatory component in lysosomes – the Vacuolar-type H+ ATPase (V-ATPase) is regulated by MTORC1[25], which was constitutively phosphorylated in the GBA1-PD mutant cells. Ratto et al.[25] demonstrated that under nutrient-deprived conditions, MTORC1 is inactive and is distributed in the cytosol, enabling the peripheral ATP6V_1_ to bind to the membrane-bound ATP6V_0_ domain, forming a functional V-ATPase complex allowing proton exchange and acidification of the lysosomes.

Under nutrient-rich conditions, MTORC1 is phosphorylated and remains on the lysosomal membrane, preventing the formation of a functional V-ATPase complex[25]. While the V-ATPase maintains lysosomal pH, disruption of the complex does not hamper autolysosome formation[26]. Total protein estimation through western blotting revealed no significant difference in the expression of ATP6V_0_D2 and ATP6V_1_A or ATP6V_1_H between the control and mutant DA neurons (Fig 4A). To examine their expression levels specifically in lysosomes, we performed a lysosomal enrichment assay and probed for pMTORC1, LAMP1, and the ATP6V_0_ and ATP6V_1_ components (Fig 4Bi). Equal amounts of cell lysate and purified lysosomes (25mg) were used to run the western blots. LAMP1 was used as a marker to verify lysosomal enrichment and GAPDH was used as a marker for cell lysates (Fig 4Bi). For quantification (Fig 4Bii), the bands in the lysosome-enriched samples were normalised to the protein levels in the lysosome-enriched samples from untreated isogenic controls. Since pMTORC1 represents the active form of MTORC1 and is found on the lysosomal membrane, we measured only pMTORC1 levels. pMTORC1 expression was increased up to two-fold, and LAMP1 expression was increased ∼1.5 fold in the GBA1-mutant DA neurons. However, ATP6V_1_A expression was reduced by ∼40% and ATP6V_1_H by ∼15% within the lysosomes fractionated from *GBA1* mutant DA neurons, while ATP6V_0_D2 levels remained unchanged between the control and mutant iPSC-DA neurons (Fig 4Bii). Furthermore, overnight treatment of *GBA1* iPSC-DA neurons with the MTORC1 inhibitor rapamycin (200nM) decreased the pMTORC1 and LAMP1 levels, while increasing ATP6V_1_A and ATP6V_1_H expression in the lysosomes and the ATP6V_0_D2 levels remained constant (Fig 4Bi b and Bii).

**Figure 4:**
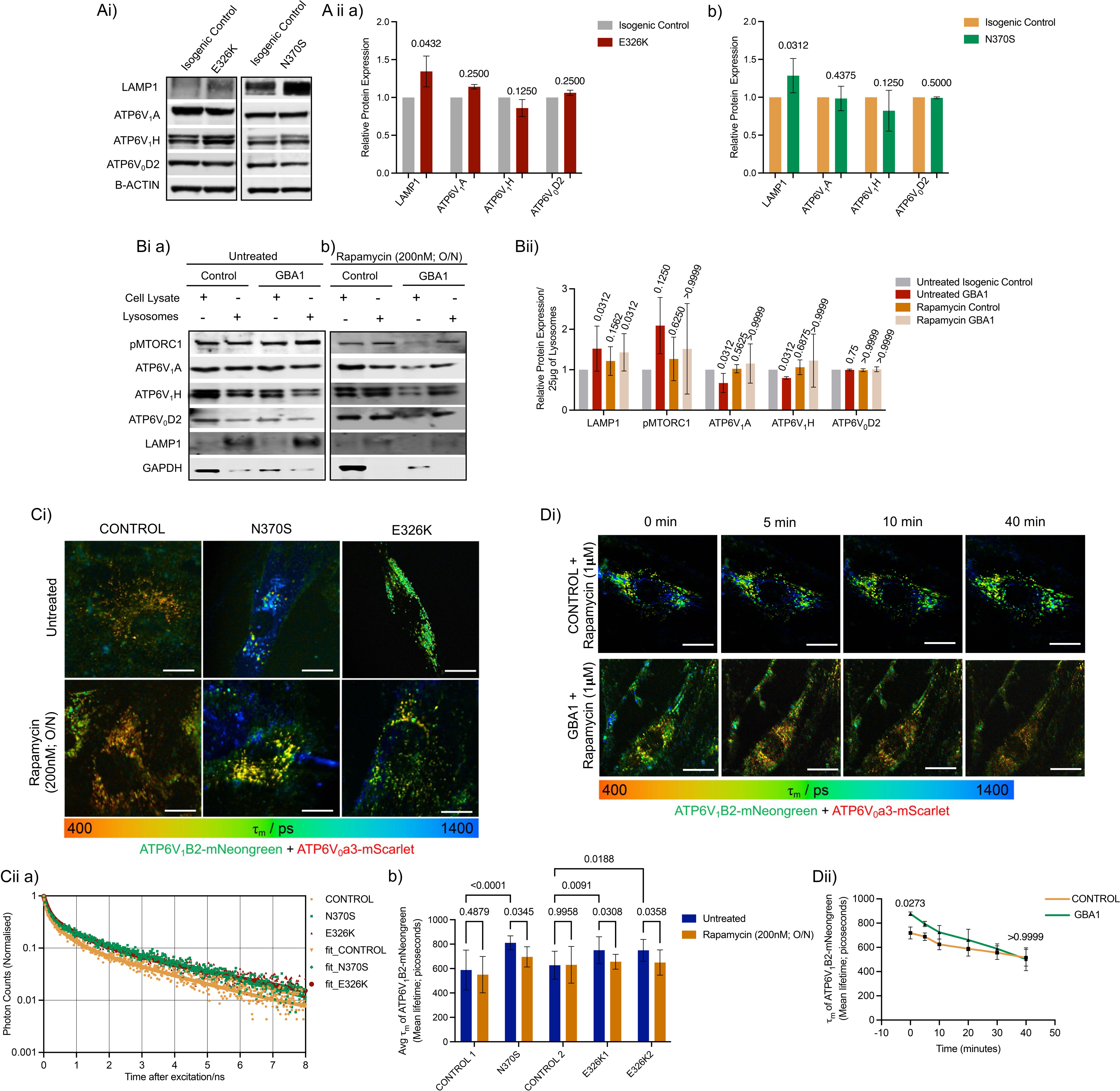
Impaired V-ATPase complex formation in GBA1-PD cells. Representative images of western blots (Ai) and quantification of protein levels in total cell lysates from isogenic control and GBA1-E326K (Aii a) and GBA1-N370S (Aii b) mutant DA neurons probed against LAMP1 and V-ATPase complex proteins. Representative immunoblotting images of cell supernatant and lysosomes separated by lysosome enrichment assay in untreated (Bi a) and 200nM rapamycin-treated (Bi b) GBA1-PD DA neurons and isogenic controls, and subsequent quantification (Bii) of V-ATPase complex proteins and pMTOR. Representative FLIM images of ATP6V_1_B2-mNeonGreen in control and GBA1-PD fibroblasts co-transfected with ATP6V_0_a3-mScarlet and treated with/without 200nM rapamycin (Ci) and subsequent quantification of mean lifetime (τ_m_) (Cii b). Cii a) Representative decay plots and exponential fit of ATP6V1B2-mNeongreen in the Control and N370S and E326K mutant fibroblasts. D i) Time point images of ATP6V_1_B2-mNeongreen in control and GBA1-PD fibroblasts co-transfected with ATP6V_0_a3-mScarlet and treated with 1 µM rapamycin and imaged over a period of 40 mins and subsequent quantification of τ_m_ shown in Dii) Data represented as mean±SD; Statistics: one-sample Wilcoxon rank test or Ordinary two-way ANOVA with Tukey’s multiple comparisons test with single pooled variance. p-values are noted on the graphs. The statistical tests, corresponding effect sizes, and confidence intervals for each graph are in supplementary table 2.

In order to assess the V-ATPase assembly, we transfected the control and *GBA1* mutant fibroblasts with ATP6V_1_B2-mNeongreen and ATP6V_0_a3-mScarlet and quantified their FRET interaction using fluorescence lifetime imaging microscopy (FLIM)[27]. The data were best fit to a biexponential decay function. The ATP6V_0_a3 construct utilises the more rapidly maturing but photophysically heterogeneous acceptor variant mScarlet-I[28]. This exhibits at least two fluorescence lifetimes, meaning the biexponential decay of the donor is an oversimplification given the likelihood of contrasting FRET rates to each acceptor species [29,30]. The results are therefore presented as the mean (amplitude weighted) fluorescence lifetime -τ_m_- which will nevertheless respond to variations in FRET without having to interpret the meaning of individual decay components and amplitudes.

τ_m_ of ATP6V_1_B2-mNeongreen averaged 617.72± 25.3 ps in control fibroblasts but increased to 760.88± 20.36 ps in N370S and 811.03± 17.04 ps in E326K mutant fibroblasts, indicating an increased distance between ATP6V1B2-mNeongreen and ATP6V0a3-mScarlet signalled by a reduction in FRET. Overnight treatment with 200nM rapamycin significantly decreased τ_m_ to 652.7± 19.69 ps and 726.05± 21.2 ps in E326K and N370S fibroblasts respectively while the τ_m_ control fibroblasts was at 590.5± 32.7 ps (Fig 4C). To validate the subcellular localisation process within a condensed timeframe, we administered 1 µM rapamycin to the cells and performed FLIM every 10 minutes for 40 minutes. The mean fluorescence lifetime of ATP6V1B2-mNeongreen exhibited a progressive decline in the *GBA1* mutant fibroblasts, decreasing from ∼850 ps at 0 min to ∼580 ps at 40 minutes, while τ_m_ in control fibroblasts showed only a small reduction, decreasing from ∼675 ps at 0 min to ∼588 ps at 40 minutes (Fig 4Dii). The FLIM-FRET data thus confirmed impaired assembly of the V -ATPase complex in *GBA1* mutant cells.

All these data are consistent with a model in which phosphorylation of MTORC1 at the lysosome membrane limits the formation of a functional V-ATPase complex in the GBA1-PD cells. Employing rapamycin as a potent MTORC1 inhibitor reduced MTORC1 activity and facilitated the formation of a functional V-ATPase complex.

### Acidification of lysosomes is sufficient to restore lysosomal, mitochondrial function and mitophagy in GBA1-PD

Our data show that *GBA1* mutations lead to significantly impaired lysosomal acidification and mitophagy and suggest that this failure of cellular homeostasis may underlie the mitochondrial dysfunction seen in GBA1-PD cells. We therefore wondered whether restoring lysosomal pH could rescue these defects in GBA1-PD cells. Since impaired V-ATPase complex formation may be a consequence of MTORC1 hyperphosphorylation, we explored the impact of treatment with rapamycin (overnight treatment with 200nM) on lysosomal and mitochondrial function. As an independent pH modulator, we employed novel poly(ethylene tetrafluorosuccinate-co-succinate) nanoparticles (NPs), which acidify lysosomes[31].

We first validated the effects of rapamycin and the nanoparticles in patient-derived fibroblasts (Fig S4). To confirm that the NPs were localised to the lysosomes, we loaded the fibroblasts with rhodamine-tagged NPs and stained the cells with Lysotracker blue DND22 (Fig S4A) which confirmed the localisation of the NPs to lysosomes. Overnight treatment with 180 µg/mL acidic NPs composed of poly(ethylene tetrafluorosuccinate-co-succinate) restored lysosomal pH in *GBA1* mutant cells to levels comparable to those of control fibroblasts as measured using the pH-sensing ratiometric probe Lysosensor Yellow/Blue DND160. Treatment with non-acidic control NPs did not significantly affect the lysosomal pH in the *GBA1* mutant fibroblasts (Fig. S4B, C). TMRM staining demonstrated that both rapamycin and the acidic NPs significantly rescued the ΔΨm in the mutant fibroblasts (Fig S4F, G). We also found that the ΔΨm was increased in control 2 and E326K fibroblasts upon control NP treatment (Fig S4G). We wondered whether the effect of control NPs on ΔΨm could be due to the succinate component in the poly(ethylene succinate) control NPs. However, treatment with 3mM Diethyl Succinate for 30 minutes did not have any effect on the ΔΨm in control and *GBA1* mutant fibroblasts (Fig S4H).

We then assessed the impact of rapamycin and acidic NPs on *GBA1* mutant iPSC-derived DA neurons (Fig 5). Treatment with 200nM rapamycin and 180 µg/mL acidic NPs restored lysosomal pH, ΔΨm, and mitophagy in the E326K and N370S iPSC DA neurons (Fig 5A, B, C). Although there was no change in GBA1 expression levels, GCase activity was increased in *GBA1* mutant DA neurons following treatment with either rapamycin or acidic NPs (Fig S4 D, E). Additionally, western blotting of OXPHOS complexes indicated that the increased mitochondrial complex IV levels in E326K DA neurons and the reduced complex I levels in the N370S DA neurons were normalised upon treatment with either rapamycin or acidic NPs. Complex III levels were also rescued upon rapamycin or acidic NPs treatment in both E326K and N370S DA neurons (Fig S4I).

**Figure 5:**
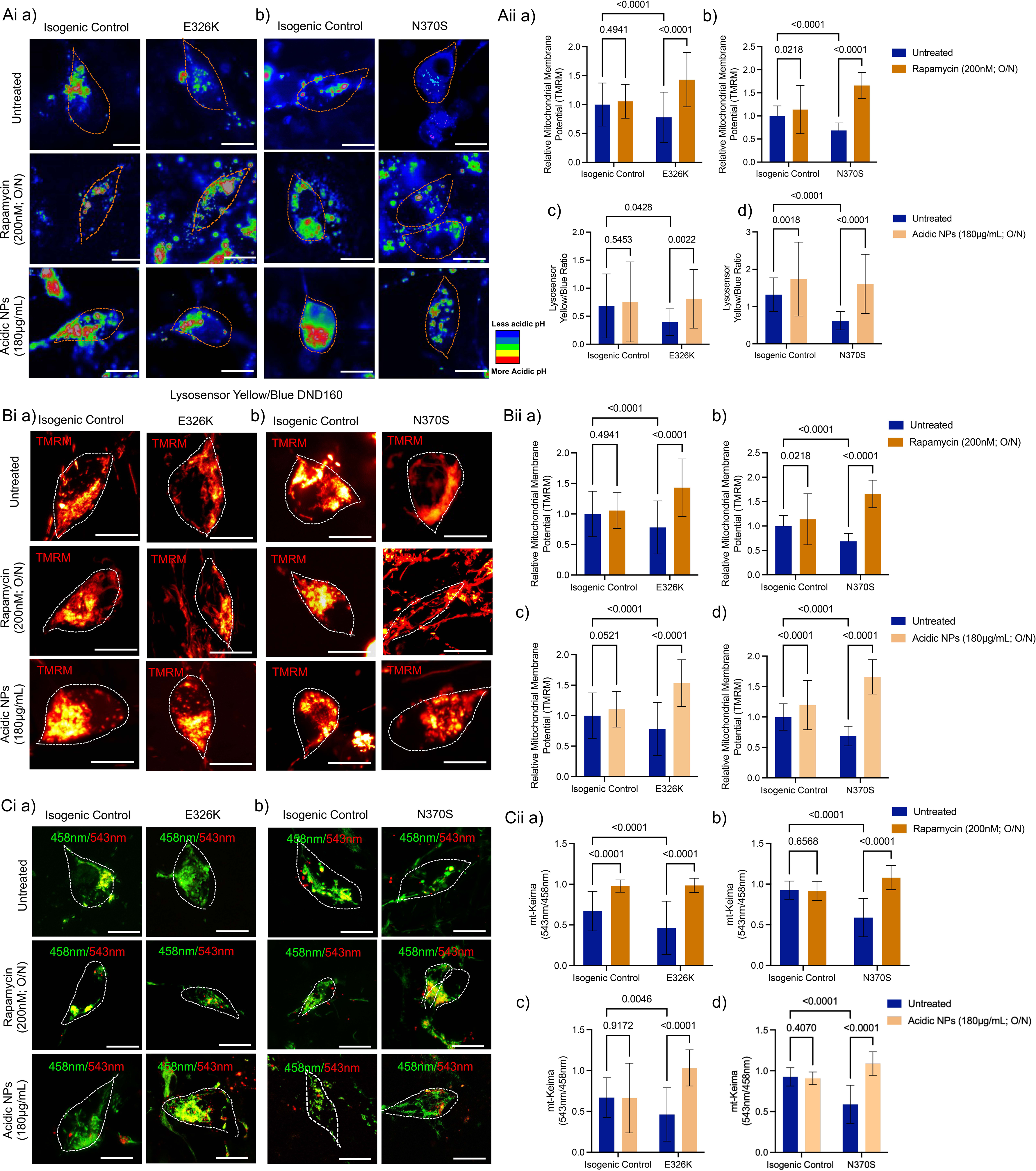
Rapamycin and acidic NPs rescue lysosomal and mitochondrial dysfunction in *GBA1* mutant DA neurons. Representative ratioed images of GBA1-E326K (Ai a) and GBA1-N370S (Ai b) DA neurons and respective isogenic controls treated overnight with 200nM rapamycin or 180μg/mL acidic NPs and stained with Lysosensor Yellow/Blue DND 160 (Ai a,b) and quantification of fluorescence ratio (Aii a-d) indicating lysosomal pH. Representative confocal images of GBA1-E326K (Bi a) and GBA1-N370S (Bi b) DA neurons and respective isogenic controls treated with 200nM rapamycin or 180μg/mL acidic NPs and stained with TMRM to measure changes in ΔΨm (Bii a-d). Representative confocal Images of GBA1-E326K (Ci a) and GBA1-N370S (Ci b) DA neurons and respective isogenic controls treated with 200nM rapamycin or 180μg/mL acidic NPs and probed with mt-Keima plasmid and imaged with excitation at 458 and 543nm and subsequent quantification of the ratio of signals at 543/458nm (Cii a-d). Data represented as mean±SD; Statistics: Ordinary two-way ANOVA with Uncorrected Fisher’s LSD with single pooled variance. p-values are noted on the graphs. The statistical tests, corresponding effect sizes, and confidence intervals for each graph are in supplementary table 2.

As rapamycin restores the assembly of the V-ATPase complex in *GBA1* mutant cells (Fig 4D-F), we asked whether the acidic NPs also function through the V-ATPase complex formation. FLIM-FRET experiments conducted on control and acidic NP-treated *GBA1* mutant fibroblasts transduced with ATP6V_1_B2-mNeongreen and ATP6V_0_a3-mScarlet plasmids demonstrated a further increase in the τ_m_ of ATP6V1B2 in both control and mutant fibroblasts treated with acidic NPs, confirming that the acidic NPs operate independently of V-ATPase (Fig S4J).

Thus, the data presented here suggest that constitutive phosphorylation of MTORC1 in GBA1-PD cells leads to impaired lysosomal pH and compromised lysosomal proteolytic function. This results in the accumulation of mitolysosomes and impaired mitophagy. The compromised mitophagy culminates in mitochondrial dysfunction in *GBA1* mutant cells. Furthermore, inhibiting MTORC1 activity or pharmacologically decreasing lysosomal pH using acidic NPs enhances lysosomal proteolytic activity, rescuing mitochondrial dysfunction in *GBA1* mutant iPSC-DA neurons and fibroblasts.

## Discussion

We report a range of abnormalities in the lysosomes and mitochondria of GBA1-PD fibroblasts and iPSC-DA neurons carrying E326K or N370S *GBA1* mutations. Given that GBA1 is a lysosomal-resident protein, we hypothesised that mitochondrial dysfunction is likely to reflect a primary lysosomal defect. We used the ratiometric probe Lysosensor Yellow/Blue DND 160 to measure lysosomal pH. When calibrated, these measurements revealed that lysosomal pH in control cells ranged from 4.74 ± 0.03 to 4.77 ± 0.02, while in mutant cells it showed a significant shift to a range of 4.89 ± 0.02 to 5.06 ± 0.03. Reports in the literature indicate that the optimal pH for lysosomal function ranges between 4.2 and 4.8, depending on the cell type [8,32]. Although the difference between control and mutant cells seems modest, the data we present suggest that this difference in mutant cells is functionally significant. In line with other studies, we observed a pH-dependent decrease in the lysosomal enzyme activity, an increase in lysosomal number, and disruption of mitophagy and mitochondrial metabolism [18,33–39].

Very few reports discuss mitochondrial abnormalities associated with GBA1-PD [35,40–43], and even fewer explore the status of mitophagy pathways mitophagy [44]. The TMRM data indicated reduced ΔΨm and also revealed mitochondrial fragmentation. EM experiments supported these findings, where mitochondria were shorter, swollen, and mitochondrial aspect ratio was reduced compared to controls. These data, along with reduced density of cristae, collectively indicate structural abnormalities in the mitochondria. A decline in ΔΨm is known to destabilise inner membrane organisation and promote both swelling and fragmentation[45], which aligns with the morphological shift toward more rounded, fragmented mitochondria observed in our analyses. We also found a significant reduction in basal respiratory rates and substrate-dependent mitochondrial respiration in the E326K and N370S DA neurons. The compromised bioenergetics and structural abnormalities collectively indicate dysfunctional mitochondria. Mitochondrial complex III levels were reduced in GBA1-E326K and N370S DA neurons compared to the respective isogenic controls, quantified in western blots. Mitochondrial complex III plays a key role in driving respiration and maintenance of membrane potential [46] and so reduced expression of complex III could account for the reduced mitochondrial membrane potential and reduced respiratory rate in GBA1-PD DA neurons. Reduced mitochondrial complex III expression has also been reported in platelets and lymphocytes of PD patients[47,48]. However, increased complex IV levels were observed in only two E326K mutant DA neurons, while one N370S line exhibited increased complex I expression. This could be due to variation between subjects. More replications and other experiments need to be performed to obtain conclusive data on mitochondrial complex levels in the E326K and N370S GBA1-PD lines. Mitochondrial dysfunction was also associated with impaired mitophagic flux in E326K and N370S GBA1-PD fibroblasts and iPSC-DA neurons.

Numerous studies have suggested mechanistic evidence for a connection between PD and *GBA1* mutations. For example, dysfunctional mitochondria in primary neurons and astrocytes were described in the *Gba1* mouse knockout model, associated with an impaired autophagy-lysosomal pathway, resulting in the accumulation of ubiquitinated proteins and α-synuclein[6]. Various studies have explored the mechanisms behind the PD pathology in *GBA1* mutant models. A recent study emphasised that decreased GCase activity in GBA1-PD midbrain dopaminergic neuronal cells leads to defective modulation of the untethering protein TBC1D15, which regulates Rab7 GTP hydrolysis for contact untethering. This impairment results in prolonged mitochondria-lysosome contacts, potentially affecting mitochondrial dynamics and function in GBA1-linked PD[42]. Baden et al. proposed an alternative function of GBA1 in the mitochondria in maintaining the integrity of complex I and support of energy metabolism[40], although in the present study we found no evidence for mitochondrial localisation of GBA1 nor impaired assembly of complex I.

Respirometry assays using isolated mitochondria revealed a marked reduction in both maximal respiratory capacity and ATPase-linked oxygen consumption in GBA1-E326K and N370S mutant DA neurons, upon stimulation with complex I (pyruvate/malate) and complex II (rotenone/succinate) substrates. The impairment across both complexes points to a global mitochondrial dysfunction, potentially arising from structural abnormalities such as fragmented mitochondria and decreased cristae density—features observed in these mutants—or as a downstream consequence of lysosomal dysfunction. Notably, lysosome-related mitochondrial impairment has also been reported in Tre-2/Bub2/Cdc16 (TBC1) domain containing Kinase (TBCK) encephaloneuronopathy, supporting a broader link between lysosomal defects and mitochondrial pathology[49].

Recent evidence suggests that the primary cause of the pathologies associated with lysosomes and mitochondria in PD is the *GBA1* mutation. There appears to be a correlation between the onset of PD and lysosomal dysfunction caused by impaired GCase activity. Dysfunctional lysosomes disrupt cellular signalling networks, metabolic processes, autophagy, and the regulation of calcium (Ca²⁺) signalling, all of which may contribute to PD pathology[7,10,19,36,38]. Furthermore, both lysosomal function and autophagy are intrinsically dependent on lysosomal acidification, which is influenced by the pH-dependence of lysosomal enzyme activity and substrate degradation, as well as the regulation of lysosomal calcium efflux[50]. Li et al. also suggest that autophagy in L444P mutant GBA1-PD neurons is impaired at two stages: the initial autophagy stage and the lysosomal degradation stage. While these reports indicate a relationship between *GBA1* mutations and autophagic, lysosomal, and mitochondrial dysfunction, a mechanistic dissection of these multi-organellar dysfunctions is lacking.

Dysregulation of the MTORC1 pathway is reported in various neurodegenerative diseases, including PD and Alzheimer’s disease[51,52]. A recent study by Chen et al. suggests that hyperphosphorylation of MTORC1 in striatal inhibitory neurons diminishes the disruption of dopamine receptor activity, alongside odour preference, in transgenic mice [53]. Liu et al. demonstrate the role of MTORC1 in dopamine dynamics and synaptic plasticity[54]. These results elucidate the additional impacts of increased phosphorylated MTORC1 on PD. Kaempferol, an MTORC1 inhibitor, has also been shown to promote autophagy and to protect DA neurons in the MPTP-induced C57BL/6J–PD mouse model [55]. All this literature suggests that hyperphosphorylated MTORC1 may play a central role in the development of PD associated with *GBA1* mutations. Our study also indicates increased MTORC1 phosphorylation in GBA1-PD fibroblasts and iPSC-DA neurons. While the link between MTORC1 hyperphosphorylation and *GBA1* mutations is unclear, it may reflect a cellular stress response to lipid substrate accumulation under *GBA1* mutant conditions[56–58].

Recent evidence underscores the significance of impaired lysosomal acidification in various disease processes. Notably, impaired lysosomal acidification is proposed to be a critical factor in the pathogenesis of multiple neurodegenerative diseases[13, 24, 27, 56]. Mutations in presenilin-1 (*PS1*) associated with familial Alzheimer’s disease have been demonstrated to inhibit the lysosomal acidification of human *PS1* mutant fibroblasts, resulting in chronic alterations in autophagy and degradation, which may be restored through lysosomal reacidification[59]. Furthermore, it has been established that individuals with Down Syndrome, who are predisposed to early-onset dementia resembling Alzheimer’s disease, exhibit increased levels of the β-cleaved carboxy-terminal fragment of the amyloid precursor protein due to the presence of an additional chromosome 21, which impairs lysosomal acidification and functionality via the inhibition of V-ATPase[17]. Collectively, these studies illuminate the connections between the impairment of lysosomal acidification in autophagic pathways and the pathogenesis of neurodegeneration, thereby presenting potential therapeutic avenues for targeting lysosomal function and pH to mitigate these debilitating diseases. Acidic NPs in the form of poly(lactic-co-glycolic acid) (PLGA) acidic NPs have been previously used to reduce lysosomal pH and disease progression in cells derived from GBA1-N370S PD patient fibroblasts and in PD mouse models[61]. Additionally, poly-succinate acidic NPs used in this study have been employed to restore lysosomal pH and rehabilitate metabolic and lysosomal functions in cases of non-alcoholic fatty liver disease[31].

As an MTORC1 inhibitor[62], rapamycin also acts as a prominent inducer of mitophagy and has been demonstrated to mitigate mitochondrial dysfunction in various disease models through inhibition of the MTORC1 pathway[63,64]. Consistent with findings from other studies, we found that rapamycin rescues mitophagy and mitochondrial function in GBA1-PD fibroblasts and DA neurons. This could either be due to the role of MTORC1 as a regulator of mitochondrial function[65,66] or a downstream effect of lysosomal acidification by MTORC1 inhibition. To establish whether impaired lysosomal acidification is the cause of these dysfunctions, we employed an independent pH modulator in the form of novel acidic poly-succinate NPs, which rectify lysosomal pH in the cells while enhancing lysosomal and mitochondrial functions in GBA1-PD fibroblasts and DA neurons. Our data confirmed the efficacy of acidic NP treatment on lysosomal pH *in vitro*. We show that the ratio of alkaline vs acidic lysosomes decreased significantly in all mutants and most controls after the acidic NP treatment, demonstrating a decrease in mean lysosomal pH. The effect on controls could be due to the alkalinisation of the lysosomes with age, which can be reversed by the acidic NP treatment in healthy cells[67]. Importantly, treatment with acidic NP successfully eliminated the differences in lysosomal pH between the *GBA1* mutants and the controls. We conclude that the restoration of lysosomal acidification in the mutant cells improved lysosomal function.

Lysosomal acidification also rescued GCase activity in the cells despite mutations in the GBA1 protein. Other studies have also demonstrated the restoration of GCase activity despite *GBA1* mutations under acidic pH conditions[68–70]. The N370S mutation results in a structurally rigid protein with reduced intra-lysosomal stability and reduced GCase activity[70], and the E326K mutation has little effect on the active catalytic site of the protein but affects the dimerisation and quaternary structure formation due to folding defects, likely leading to a mild reduction in GCase activity at higher pH levels[68]. GCase activity depends on protonation of amino acids at the active sites and due to interaction of the protein with activators like saposin C[71], which is affected due to the mutation[68,72]. Restoring the acidic pH in the lysosomes helps restore functional GCase activity despite the mutations possibly by stabilising protein structure, improving folding and optimising the active-site ionisation[68,73]

To explore whether altered lysosomal pH is related to mitochondrial dysfunction in GBA1-PD, we evaluated ΔΨm, redox levels, and oxidative phosphorylation (OXPHOS) protein levels following treatment with NPs. Our results reveal a significant rescue of ΔΨm among GBA-PD mutants after acidic NP treatment, indicating improved mitochondrial function. When compared to untreated healthy controls, the difference in ΔΨm was no longer statistically significant in the mutants following treatment with acidic NPs. It is noteworthy that some control samples also exhibited alterations due to acidic NP treatment, resulting in a small but significant increase in ΔΨm. We also found a significant increase in ΔΨm in control 2 and E326K fibroblasts upon control NP treatment. Since the NPs are composed of poly(ethylene-succinate), we suspected this could be a substrate-mediated (succinate) effect in control NP treated conditions. However, 30-minute treatment with 3mM Diethyl Succinate did not alter the ΔΨm in the control or mutant fibroblasts. Acidic NP treatment also rescued the differential mitochondrial complex levels observed in the GBA1-E326K and N370S DA neurons. Rescuing lysosomal pH also improved mitophagy in the GBA1-PD fibroblasts and iPSC-DA neurons.

## Conclusions

We have explored the relationship between lysosome biology, pH, mitophagy and mitochondria in cells carrying PD-associated mutations of *GBA1*. A detailed examination of the pathophysiology of these cells reveals a range of abnormalities in the lysosomes, including reduced GCase activity, an increased lysosomal number, impaired lysosomal acidification, and impaired pH-dependent lysosomal proteolytic activity. Mitochondrial dysfunction is reflected by reduced ΔΨm, swollen, fragmented mitochondria, decreased cristae density, and the accumulation of abnormal mitochondria, along with a reduced oxygen consumption rate. We also found a decrease in mitophagy and an accumulation of autolysosomes in the GBA1-PD cells. We show that increased pMTORC1 on the lysosomal membrane, possibly due to nutrient accumulation, hinders the formation of a functional V-ATPase complex in GBA1-PD cells, thereby compromising lysosomal acidification. We found that inhibiting MTORC1 using rapamycin rescues functional V-ATPase formation and lysosomal pH in the GBA1-PD neurons, along with mitophagy and mitochondrial function. Independent modulation of lysosomal pH using acidic poly-succinate NPs, restored lysosomal pH, rescued mitophagy and restored mitochondrial function in the GBA1-PD fibroblasts and iPSC-DA neurons (Fig 6). Since rapamycin has broad effects, the comparison to NPs is valuable, adding a translational value for future applications. These studies identify the underlying mechanism causing impaired lysosomal pH regulation associated with PD-related GBA1 mutations and provide evidence that restoring lysosomal pH is sufficient to rescue lysosomal function, to restore mitophagy and therefore to rescue mitochondrial function in both GBA1-PD patient-derived fibroblasts and iPSC-DA neurons. We highlight that lysosomal pH correction is a pivotal, unifying therapeutic target in GBA1-PD.

**Figure 6:**
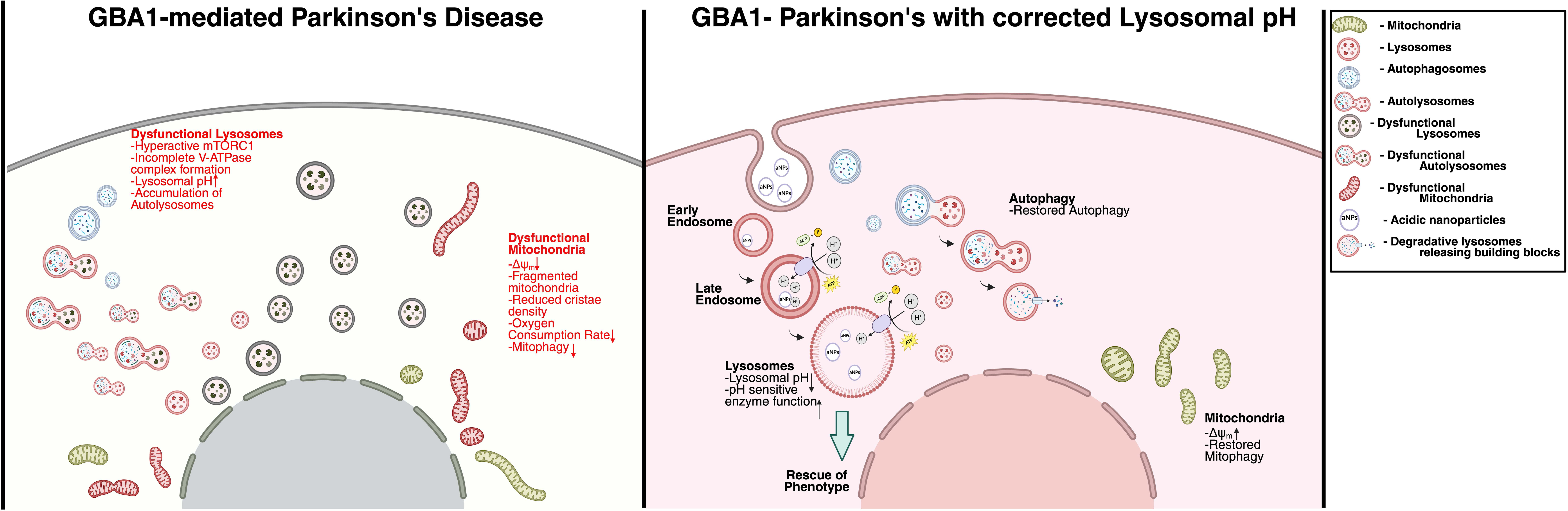
Cartoon to illustrate the proposed pathophysiological cascade in GBA1-related Parkinson’s Disease and how the phenotype can be corrected by targeting lysosomal pH. GBA1-PD cells exhibit a range of abnormalities in lysosomal biology, including reduced GCase activity, increased lysosomal numbers, elevated lysosomal pH, and decreased pH-dependent proteolytic activity. Mitochondrial dysfunction is evidenced by a decreased ΔΨm, decreased oxygen consumption rate, swollen mitochondria, reduced cristae density, and the accumulation of abnormal mitochondria. Mitophagy is suppressed and autolysosomes accumulate within the GBA1-PD cells. Restored acidification of lysosomes with rapamycin or acidic poly-succinate NPs enhances lysosomal function and improves mitophagy, thereby rescuing mitochondrial dysfunction in GBA1-PD fibroblasts and lysosomes.

---

**Supplementary Figure 1:**
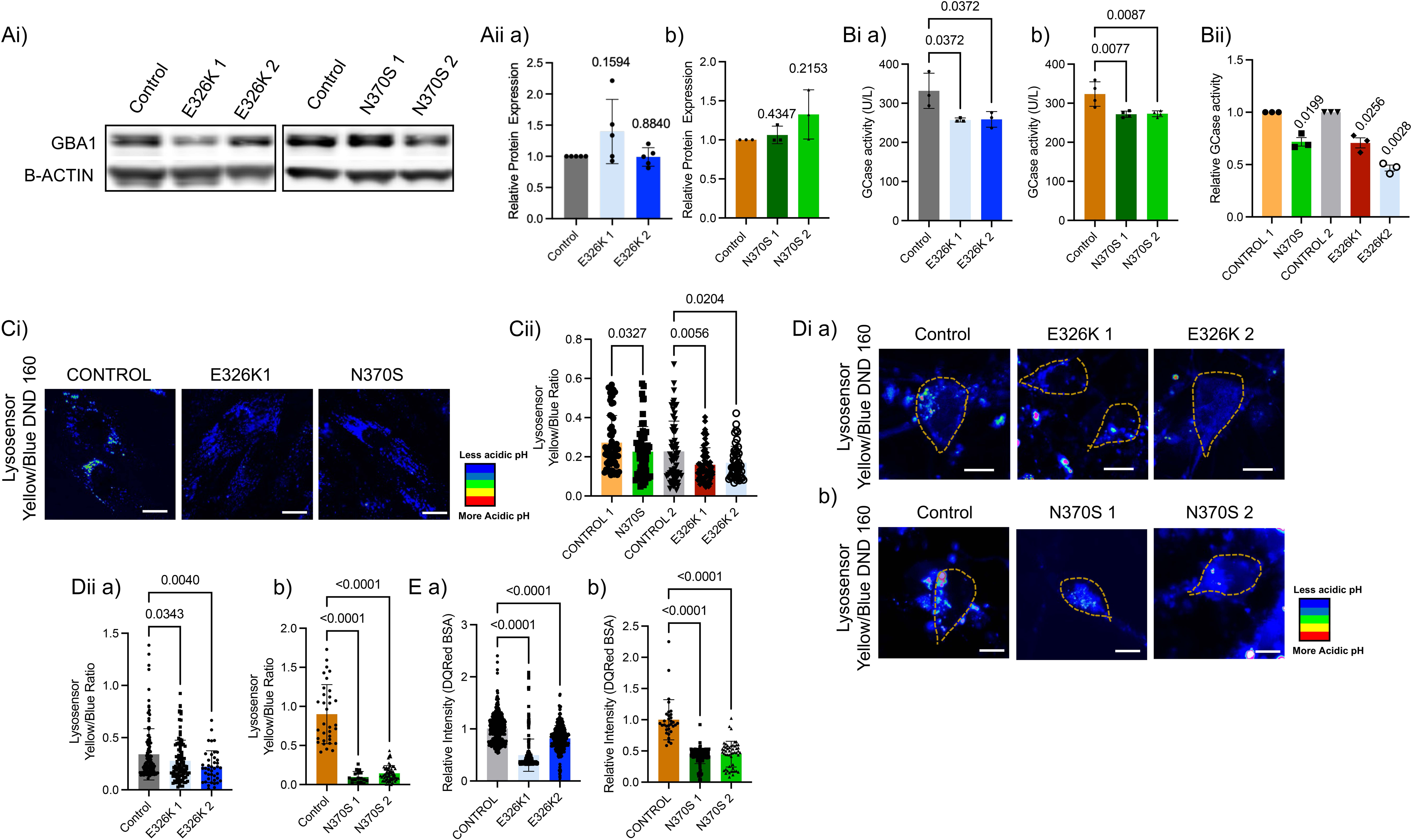
Lysosomal abnormalities in GBA1-PD DA neurons and fibroblasts. GBA1 protein expression (Ai) and subsequent quantification of GBA1 levels in control and GBA1-E326K (Aii a) and N370S (Aii b) DA neurons. GCase activity in GBA1-E326K (Bi a) and GBA1-N370S (Bi b) DA neurons and fibroblasts (Bii). Representative ratioed images of GBA1-E326K and GBA1-N370S fibroblasts (Ci) and DA neurons (Di a,b) stained with Lysosensor Yellow/Blue DND 160, subsequent quantification of fluorescence ratio measuring lysosomal pH (Cii, D ii a,b). Histogram representing quantification of DQ-Red BSA intensity in GBA1-E326K (E a) and GBA1-N370S (E b) and control DA neurons indicating lysosomal proteolytic activity. Data represented as mean±SD; one-sample Wilcoxon rank t-test or One-way ANOVA with Kruskal-Wallis and Dunn’s multiple comparisons test or Holm-Šídák’s multiple comparisons test. p-values are noted on the graphs. The statistical tests, corresponding effect sizes, and confidence intervals for each graph are in Supplementary Table 2

**Supplementary Figure 2:**
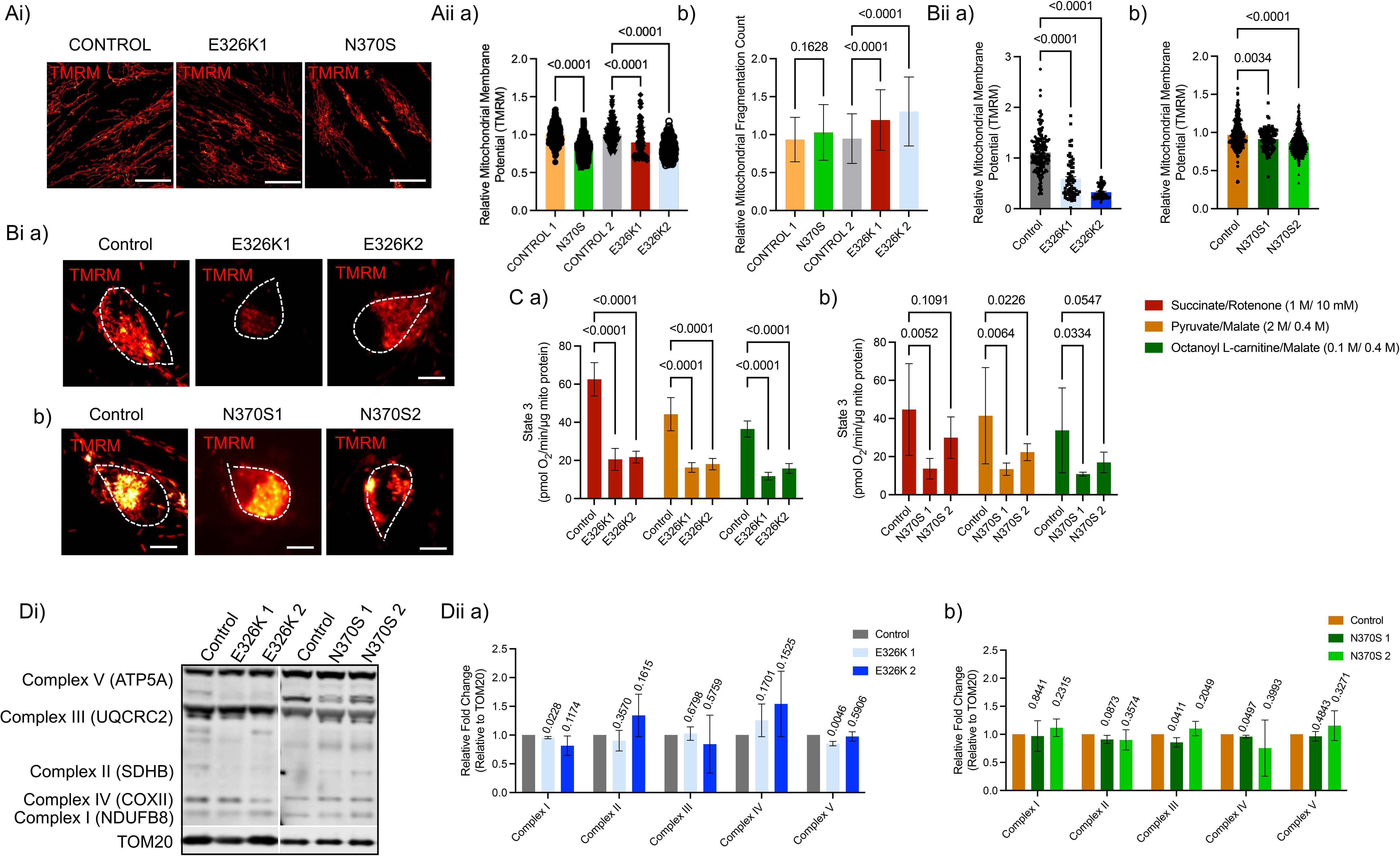
Mitochondrial dysfunction in GBA1-PD DA neurons and fibroblasts. Representative images of GBA1-E326K and GBA1-N370S fibroblasts (Ai) and DA neurons (Bi a,b) along with controls, stained with TMRM to measure changes in ΔΨm (Aii a, Bii a,b) and the mitochondrial fragmentation count (Aii b). Histograms representing State 3 (ATP Production) in mitochondria isolated from GBA1-E326K (C a) and GBA1-N370S (C b) DA neurons and controls, as measured by Seahorse assay. Representative images of western blots probed for OXPHOS complex proteins (Di) and subsequent quantification of protein expression levels in GBA1 E326K (Dii a) and N370S(Dii b) DA neurons. Data represented as mean±SD; Statistics: one-sample Wilcoxon rank t-test or One-way ANOVA with Kruskal-Wallis and Dunn’s, Holm-Šidák or Šídák’s multiple comparisons test or Ordinary two-way ANOVA with Tukey’s multiple comparisons test with single pooled variance. p-values are noted on the graphs. The statistical tests, corresponding effect sizes, and confidence intervals for each graph are in Supplementary Table 2

**Supplementary Figure 3:**
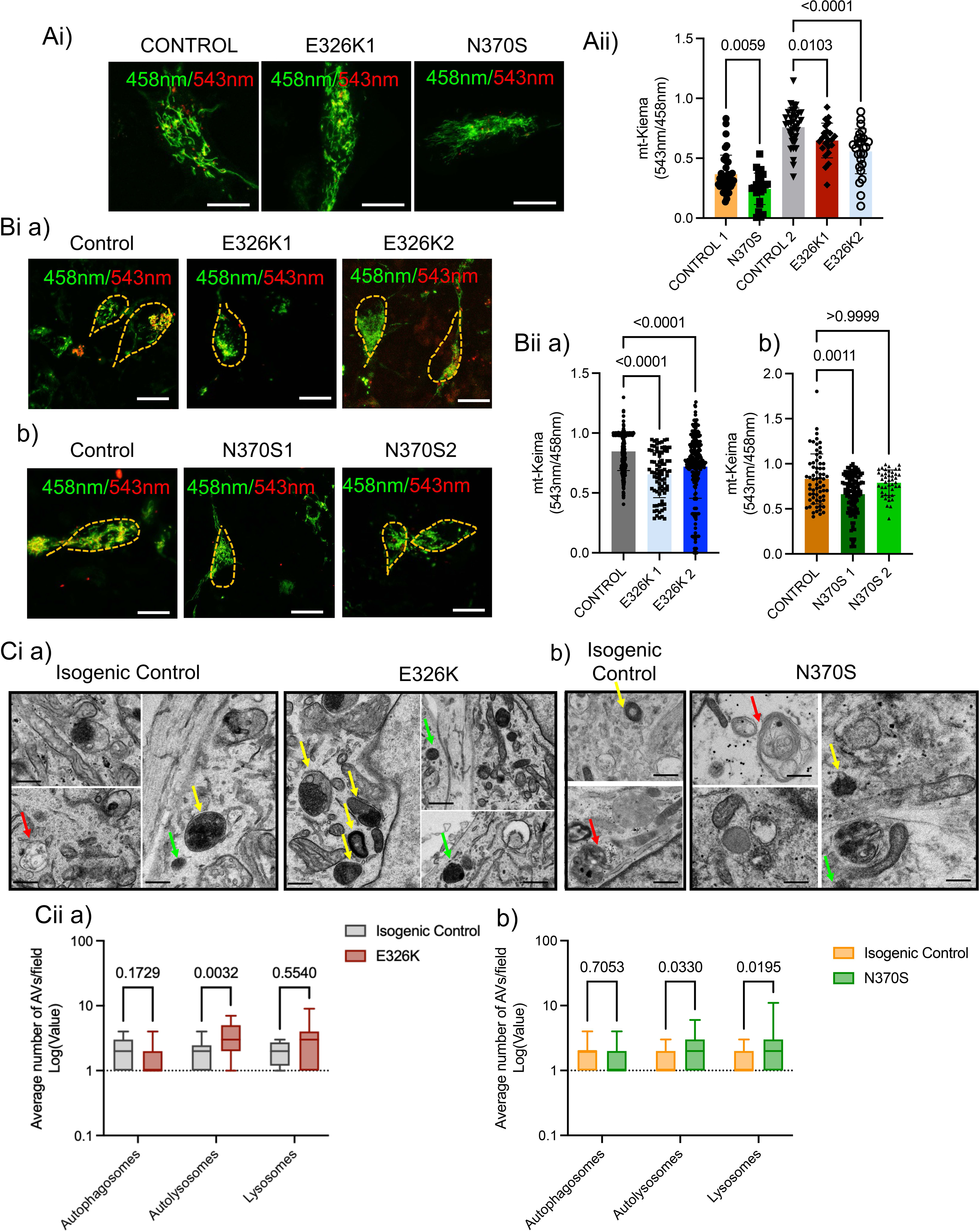
Mitophagy defects in GBA1-PD DA neurons and fibroblasts. Representative ratioed images of control and GBA1-E326K and GBA1-N370S fibroblasts (Ai) and DA neurons (Bi a,b) transduced with mt-Keima plasmid and imaged imaged with excitation at 458 and 543nm and subsequent quantification of the ratio ratio of signals at 543/458nm respectively (Aii, Bii a,b). Representative TEM images of GBA1-E326K (Ci a) and GBA1-N370S (Ci b) DA neurons and respective isogenic controls depicting autophagic vesicles and (Cii a,b) Box-and-whisker plots showing the minimum, maximum, median, and interquartile range of the log(1+Y) values. Yellow Arrows indicate autolysosomes, red arrows indicate autophagosomes, and green arrows indicate lysosomes in the cells. Data represented as mean±SD; Statistics: One-way ANOVA with Kruskal-Wallis and Dunn’s multiple comparisons test or Holm-Šidák multiple comparisons test or Ordinary two-way ANOVA with Šidák multiple comparisons test. p-values are noted on the graphs. The statistical tests, corresponding effect sizes, and confidence intervals for each graph are in Supplementary Table 2.

**Supplementary Figure 4:**
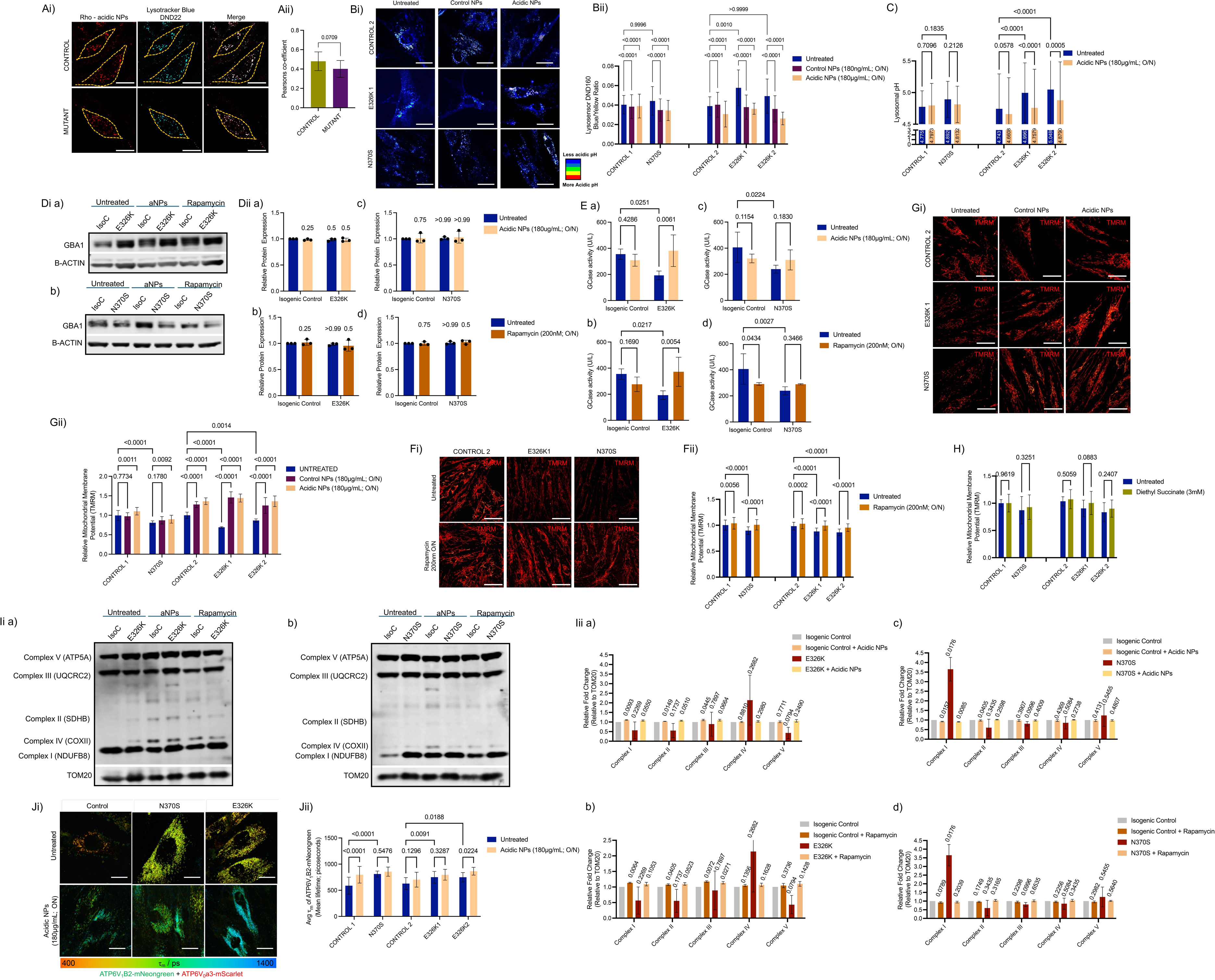
Effect of acidic NPs and rapamycin on GBA1 – PD fibroblasts and DA neurons. Representative Confocal images (Ai) of control and mutant fibroblasts treated with Rhodamine-tagged acidic NPs (Rhod-acidic NPs) and stained with Lysotracker Blue and subsequent quantification of colocalisation coefficient (Aii). Representative images of control and GBA1-E326K and GBA1-N370S fibroblasts treated with control and acidic NPs (Bi) and stained with the ratiometric Lysosensor Yellow/Blue DND160 subsequent quantification of fluorescence ratio (Bi) to measure lysosomal pH and. C) Calibration of lysosomal pH in *GBA1* mutant fibroblasts treated with and without acidic nanoparticles. Representative images of western blots and quantification of GBA1 protein levels in GBA1-E326K (Di a; Dii a,b) and GBA1-N370S (Di b; Dii c,d) DA neurons and respective isogenic controls treated with acidic NPs or rapamycin and probed for GBA1 and B-ACTIN. GCase activity in GBA1-E326K and GBA1-N370S DA neurons and respective isogenic controls treated with Acidic NPs (E a, c)/rapamycin (E b, d). Images representing rapamycin treated (Fi) and Acidic NPs treated (Gi) control and GBA1-E326K and GBA1-N370S fibroblasts and stained with TMRM and subsequent quantification of ΔΨm (Fii and Gii respectively). Histogram representing quantification of mitochondrial membrane potential in *GBA1* mutant and control fibroblasts upon 3mM Diethyl succinate treatment for 30 minutes (H). Representative images of western blots (Ii a,b) and subsequent quantification of OXPHOS complex proteins in GBA1 E326K and N370S DA neurons and respective isogenic controls treated with Acidic NPs (Iii a,b) and rapamycin (Iii c,d). Representative FLIM images of ATP6V_1_B2-mNeongreen in Control and GBA1-PD fibroblasts co-transfected with ATP6V_0_a3-mScarlet and treated with/without acidic NPs (Ji) and subsequent quantification of mean lifetime (τ_m_) of ATP6V_1_B2 (Jii). Data represented as mean±SD; Mann-Whitney’s test, one-sample Wilcoxon rank t-test or Ordinary two-way ANOVA with Tukey’s multiple comparisons test or Uncorrected Fisher’s LSD with single pooled variance. p-values are noted on the graphs. The statistical tests, corresponding effect sizes, and confidence intervals for each graph are in Supplementary Table 2

## Methodology

### Cell lines and ethics committee approval

The human iPSC (hiPSC) and fibroblast lines used in the study, along with their sources, are detailed in Table 1. Fibroblasts were cultured in DMEM (1X) + GlutaMAX™ basal media (Gibco 2206106), 10% (v/v) Foetal Bovine Serum (Gibco 10270098), and 1% antibiotic-antimycotic (Gibco 15240096), and incubated at 37 °C with 5% CO_2_. The media was changed every 72 hours, and the cells were passaged upon reaching 80% confluency using 0.25% Trypsin-EDTA (Gibco 25200056). Passages between 9 and 15 were used for experimental purposes. Human GBA1 patient-derived iPSC lines were cultured under feeder-free conditions on Geltrex-coated 6-well plates, using mTeSR Plus medium and were passaged every 4-5 days with a 0.5 mM EDTA solution or upon reaching 80% confluency. Passage numbers between 20 and 30 were employed for differentiation purposes.

### Human pluripotent stem cell-derived DA neurons

DA neurons were generated from iPSCs using a step-by-step differentiation protocol, as previously described[74]. Briefly, 90-95% confluent hiPSCs were cultured on Geltrex-coated plates and maintained in N2B27 medium, which is a 1:1 mix of Neurobasal (Gibco, 21103049) supplemented with 1X B27 (Gibco, 17504044), Glutamax (Gibco, 35050-038), and 0.5X Anti-anti (Gibco, 15240096), and DMEM-F12 (Gibco, 10565018) supplemented with 1X N2 supplement (Gibco, 17502048), 1X MEM Non-Essential Amino Acids (Gibco, 11140-050), 0.5X Anti-anti (Gibco, 15420096), 50 μM β-mercaptoethanol (Gibco, 21985-023), and human Insulin Solution (Sigma, I9278). Daily media changes were conducted for the first 14 days. On day 0, the media was supplemented with 5 μM SB431542 (Tocris Bioscience, 1614/10), 2 μM Dorsomorphin (Tocris Bioscience, 3093/10), and 1 μM CHIR99021 until day 2. From day 2 to day 7, 1 μM Purmorphamine (Merck Millipore, SML0868) was added to the mix. From day 8 to day 14, CHIR99021 and SB431542 were removed from the medium, and the cells were maintained with Dorsomorphin and Purmorphamine in N2B27. Cells were dissociated with 1 mg/mL Dispase (Gibco, 17105041) on days 4 and 14. After patterning for 14 days, the midbrain DA neuronal precursor cells (NPCs) were maintained in N2B27 medium until day 18. On day 19, the cells were dissociated with Accutase (Gibco, A1110501) and plated onto Geltrex-coated plates at a density of 2 × 10^5 cells/cm² and terminally differentiated with N2B27 medium supplemented with Compound E (Enzo Lifesciences, ALX-270-415-C250) and 10 µM Rho-kinase inhibitor, Y-27632 dihydrochloride (Tocris, 1254) from day 20 until day 70-75, with media changes twice a week. DA neurons between days 70 and 75 were used for downstream experiments.

### Nanoparticle, rapamycin and diethyl succinate treatment

Control and acidic NPs, composed of poly(ethylene-succinate) and poly(ethylene tetrafluorosuccinate-co-succinate) respectively, were obtained from Prof. Mark Grinstaff and Prof. Orian Shirihai. The synthesis of the nanoparticles is described in [31]. The nanoparticles were stored at -80°C in aliquots and thawed fresh prior to treatment. Aliquots of 75 mg/mL nanoparticles were thawed to room temperature, vortexed thoroughly, and added to the cells at a concentration of 180 µg/mL in fresh culture medium for 12-16 hours before the experiment. A stock solution of 200 µM rapamycin (Sigma, R0395) was prepared in DMSO and sterile filtered before being stored as aliquots at -20°C. This stock solution was diluted to 200 nM in medium before being added to the cells. The cells were treated with 200 nM rapamycin for 12-16 hours before proceeding with the experiments unless mentioned otherwise. Diethyl Succinate (Merck, 112402) was diluted to 3 mM in culture media and sterile-filtered and added onto cells 30 minutes prior to measuring the ΔΨm using TMRM.

### Generation of lentivirus and transduction

pLV-Ef1a-3xHA-ATP6V1B2-mNeonGreen and pLV-Ef1a-3xFlag-ATP6Voa3-mScarlet plasmids were kindly provided by Dr. Wilhelm Palm (EPR 462 and EPR 475, European Plasmid Repository). pHAGE-mt-mKeima was a generous gift from Prof. Richard Youle (131626, Addgene). Lentiviral plasmids were packaged using psPAX2 and pMD2.G in HEK293T cells with Xtremegene-HP transfection reagent (Merck, 6366236001), according to the manufacturer’s instructions. The media supernatant containing viruses was collected 48 and 72 hours post-transfection and concentrated using Lenti-X transfection reagent (Takara, 631231) with overnight incubation, followed by centrifugation at 2000 rpm for 1 hour at 4°C. The pellet was then resuspended in DMEM-F12 medium and stored at -80°C for further use. 72-96 hours prior to imaging, fibroblasts or DA neurons were treated with lentiviral particles. 24 hours post-treatment, the media was replaced with regular culture medium and incubated for another 48 hours before imaging.

### Confocal imaging

DMEM-F12 medium (Gibco, 21041025) or BrainPhys medium (Stem Cell Technologies, 05796) served as a recording buffer for live cell imaging of fibroblasts and DA neurons. All images were acquired using a Carl Zeiss LSM 880 Confocal Laser Scanning microscope with Zen Black Software and a 63×/1.40 oil immersion lens at 37°C. Fixed and stained fibroblasts were also imaged with the same equipment.

### Lysosensor Yellow/Blue DND 160

The cells were washed with DPBS and incubated for 4 minutes in the recording buffer containing 3 μM Lysosensor Yellow/Blue DND 160 (Invitrogen, L7545) at 37°C. After incubation, the dye was washed twice with PBS and imaged in fresh recording buffer using the UV laser at 355 nm. The dual emission maxima of the dye (440 nm/540 nm) were detected using spectral scanning (lambda scan mode) on the Zeiss LSM 880. Images were subsequently linearly unmixed employing the ZEN Black software to separate the 440 nm and 540 nm emission images. The ratio of 440/540 nm was then calculated using the Ratio Plus plugin on ImageJ/Fiji software to determine the lysosomal acidity of fibroblasts and DA neurons

### Lysosomal pH measurement

The lysosomal pH was measured using Lysosensor Yellow/Blue DND-160 (L7545, Invitrogen). Fibroblasts cultured on fluorodishes were treated with pH calibration buffer containing Monensin (Sigma, M5273) and Nigericin (Sigma, N7143), with a pH ranging from 3.5 to 7.0 and stained with Lysosensor Yellow/Blue DND 160. The cells were incubated with 2 µM for 5 minutes at room temperature (RT) and imaged as previously mentioned. The 440/540 fluorescence ratio was measured for each calibration buffer. A lysosomal pH calibration curve was established by correlating the 440/540 nm ratio with the corresponding pH values using ImageJ/Fiji software.

### Measurement of lysosomal proteolytic activity by DQ Red BSA

The DQ-Red BSA trafficking assay dye (Invitrogen, D12051) was utilised to examine lysosomal proteolytic activity. DA neurons were washed with DPBS, and pre-warmed medium containing DQ-Red BSA was added to the fluorodishes. The dishes were incubated at 37°C for three hours. Following the incubation period, the cells were washed with DPBS and replaced with the recording buffer prior to imaging. Cells stained with DQ-Red BSA were excited using a 561 nm Argon laser, and the emitted fluorescence was collected within the 564–740 nm range. The fluorescence intensity was quantified per cell using ImageJ/Fiji software with consistent threshold settings across all samples.

### Measurement of ΔΨm using TMRM

Fibroblast or DA neurons were washed with DPBS and incubated in recording buffer containing 20 nM TMRM (Thermofisher Scientific, T668) at 37°C for 30 minutes. Following incubation, fresh recording buffer containing 20 nM TMRM was added before proceeding with imaging. The cells were excited for TMRM fluorescence at 561 nm, and images were acquired as Z-stacks. Maximum intensity projection images were then utilised to quantify the fluorescence intensity using ImageJ/Fiji software with consistent threshold settings across all samples. The TMRM images were also used to analyse the mitochondrial morphology [75].

### Measurement of Lipofuscin levels

Lipofuscins are excited at 355 nm and fluoresce across a range from 480 to 700 nm[20]. Fibroblasts were washed with the recording buffer and imaged using UV illumination at 355 nm, with images obtained through spectral scanning. The images were subsequently linearly unmixed using ZEN Black software to separate the 460 nm and 480 nm emission images, thus distinguishing NAD(P)H from lipofuscins. The spectrally unmixed images at 480 nm were then utilised to calculate lipofuscin density per cell using ImageJ/Fiji software.

### Colocalisation of the Rhodamine-B labelled acidic NPs with the lysosomes

Rhodamine B-labelled acidic NPs were treated for 12-16 hours prior to imaging the fibroblasts. On the imaging day, cells were washed with the recording buffer and incubated with 50 nM LysoTracker Blue DND-22 dye (Invitrogen, L7525) for 2 hours at 37°C. Following the incubation, the cells were washed with the recording buffer and imaged in fresh buffer. Images were acquired using sequential excitation (405 nm for LysoTracker Blue and 561 nm for Rhodamine) and emission ranges set to 420–480 nm and 570–620 nm, respectively. Colocalisation Pearson’s coefficient was quantified using ImageJ/Fiji software with consistent threshold settings across all samples.

### Measurement of mitophagy using mt-Keima reporter

Measurement of mitophagy was performed as previously described[76]. Untreated fibroblasts and DA neurons were tranduced with lentiviral mt-Keima particles 72 hrs prior to imaging. 12-16 hours before imaging, the cells were treated with acidic NPs or rapamycin. The cells were imaged using two sequential excitation wavelengths (458 nm for green fluorescence and 561 nm for red fluorescence) with an emission range of 570–695 nm. The laser power was set at the minimum output to allow the clear visualisation of the mt-Keima signal. The ratio of the high F_543_:F_458_ ratio values were generated using the Ratio Plus plugin in ImageJ/Fiji and was used as an index of mitophagy.

### Fluorescence lifetime imaging microscopy (FLIM) quantification of ATP6V_1_B2-mNeonGreen

Fluorescence lifetime imaging was conducted using single-photon excitation on a multimodal time-resolved fluorescence microscope as previously described[77]. This setup encompassed an 80 MHz, near-infrared, femtosecond excitation source (Insight X3, Spectra Physics, Crewe, UK), a second harmonic generation unit (Harmonixx SHG, APE, Berlin, Germany), a laser scanning unit (DCS-120, Becker & Hickl, Berlin, Germany), an inverted microscope (Axio Observer 7, Zeiss, Cambridge, UK) featuring a high numerical aperture objective (Plan-Apochromat 63x/1.4 Oil M27, Zeiss, Cambridge, UK), an ultrafast hybrid detector (HPM-100-07, Becker & Hickl, Berlin, Germany), and time-correlated single photon counting (TCSPC) electronics (SPC-180NX, Becker & Hickl, Berlin, Germany). Images were acquired using 473 nm excitation to minimise the ratio of acceptor to donor excitation, along with 500–540 nm emission filtering to isolate fluorescence from the mNeonGreen donor. Photon counts were acquired for two minutes and histogrammed at 14.6ps intervals. Curve fitting analysis was performed in SPCImage (Becker & Hickl, Berlin, Germany).

### Lysosomal enrichment assay

A lysosomal enrichment assay was conducted using the Lysosome Enrichment Kit for Tissues and Cultured Cells (Thermo Scientific, 89839) according to the manufacturer’s instructions. Briefly, the DA neurons were pelleted and lysed with lysosome enrichment reagent A, which contained 1X protease and phosphatase inhibitors. The solution was then sonicated on ice, applying 12 bursts at 9W of power (Thermo Scientific) and mixed with lysosome enrichment reagent B. The solution was subsequently centrifuged at 500g for 10 minutes. 200 µL of supernatant was then aliquoted for use as cell supernatant for western blotting. Lysosomes were isolated from the remaining solution through gradient centrifugation. The protein concentrations of the lysosomes and cell supernatant were then quantified using a BCA assay and proceeded for western blotting as discussed below.

### SDS-PAGE and immunoblotting

Fibroblasts and DA neurons were washed with ice-cold PBS, followed by the addition of 150 μl of ice-cold RIPA lysis buffer (Sigma-Aldrich, R0278), which was supplemented with protease inhibitors (Roche 4693116001), PMSF (Sigma, 93482), and phosphatase inhibitors (Roche 4906837001). Cells were scraped using a plastic scraper, and the lysates were transferred to 1.5 ml tubes. The lysates were rotated at 4°C for 30 minutes and sonicated (3 cycles, 3 seconds each at 40% amplitude, with 5-minute intervals). Samples were centrifuged at 16,000 g for 30 minutes at 4°C, and the supernatant was collected. Protein concentration was determined using a BCA assay kit (Thermo Scientific, 23227). A total of 20-30 μg of protein was diluted with RIPA buffer and mixed with NuPAGE 4X sample buffer (Invitrogen, NP0007). The samples were heated at 95°C for 5 minutes (for OXPHOS proteins, the lysates were heated at 45°C for 5 minutes). Proteins were separated on 4–12% NuPAGE Bis-Tris polyacrylamide gels (Invitrogen, NP0335) immersed in MOPS running buffer (Invitrogen, NP0001). Proteins were transferred to PVDF membranes (Millipore, IPFL00010) activated in methanol using a wet transfer system. Membranes were blocked in Superblock blocking buffer (Invitrogen, 37545) for 1 hour at room temperature. The blots were cut where necessary before incubating with primary antibodies, which were diluted in 1X blocking buffer, and incubated with the membranes overnight at 4°C. After three 10-minute washes in TBST, the membranes were incubated with secondary antibodies (Li-COR Biosciences; 1:10000; IRDye® 680RD Goat anti-Mouse IgG, 926-68070; IRDye® 800CW Goat anti-Rabbit IgG, 926-32211) diluted in 1% BSA/TBST for 1 hour at room temperature. The membranes were washed three additional times with TBST. Fluorescent signals were developed using a LiCOR Odyssey CLx system. The details of the antibodies are listed in Table 2.

**Table 2.** List of antibodies used in the study.

| **Name** | **Catalogue**<br>**no.** | **Host** | **Company** | **Application** | **Concentration** |
| --- | --- | --- | --- | --- | --- |
| ATP6V0D2 | AB194557 | Mouse | Abcam | Western Blotting | 1:1000 |
| LAMP1 (H4A3) | sc-20011 | Mouse | Santa Cruz | Western Blotting,<br>Immunofluorescence | 1:500 |
| GBA (c-term) | 64171 | Rabbit | Sigma | Western Blotting | 1:700 |
| ATP6V1H | AB187706 | Rabbit | Abcam | Western Blotting | 1:1000 |
| ATP6V1A | 199326 | Rabbit | Abcam | Western Blotting | 1:1000 |
| pMTOR | 55365 | Rabbit | Abcam | Western Blotting | 1:1000 |
| MTOR | 45175 | Mouse | Cell Signalling | Western Blotting | 1:1000 |
| ATG5 (D5F5U) | 12994T | Rabbit | Cell Signalling | Western Blotting | 1:1000 |
| P62 | 610833 | Mouse | BD Biosystems | Western Blotting | 1:1000 |
| LC3 | L7543 | Rabbit | Sigma | Western Blotting | 1:1000 |
| B-ACTIN | ab8226 | Mouse | Cell Signalling | Western Blotting | 1:5000 |
| TOM20 | ab186735 | Rabbit | Abcam | Western Blotting | 1:1000 |
| LC3 | PM036 | Rabbit | MBL Biosystems | Immunofluorescence | 1:500 |
| Citrate<br>Synthetase | ab96600 | Rabbit | Abcam | Immunofluorescence | 1:200 |
| OXPHOS<br>Cocktail | 45-8199 | Mouse | Thermoscientific | Western Blotting | 1:1000 |
| Alexa Fluor 488 | A-11008 | Goat | Thermoscientific | Immunofluorescence | 1:1000 |
| Alexa Fluor 594 | A-11012 | Goat | Thermoscientific | Immunofluorescence | 1:1000 |
| Alexa Fluor 647 | A-21235 | Goat | Thermoscientific | Immunofluorescence | 1:1000 |
| IRDye® 680RD<br>Goat anti-Mouse | 926-68070 | Goat | Li-COR<br>Biosciences | Western Blotting | 1:10000 |
| IRDye® 800CW<br>Goat anti-Rabbit<br>IgG | 926-32211 | Goat | Li-COR<br>Biosciences | Western Blotting | 1:10000 |

### Immunostaining

Fibroblasts were grown on coverslips, treated, and fixed with 4% (w/v) paraformaldehyde. After fixation, the cells were permeabilised with 50 µg/ml digitonin in PBS for 10 minutes. The cells were then washed, blocked with 3% BSA, and incubated with the following primary antibodies: citrate synthase, LAMP1, and LC3 in 3% BSA for 1 hour at room temperature, followed by incubation with Alexa Fluor 488/594/647-conjugated secondary antibodies for 1 hour at room temperature. Coverslips were mounted on glass slides, and images were acquired using appropriate excitation and emission filters to capture fluorescent signals. Colocalisation Pearson’s coefficient (Mito vs LC3 and Mito vs LAMP1), lysosomal number (LAMP1 particles/cell), and autophagosome number (LC3 particles/cell) were quantified using ImageJ/Fiji with consistent threshold settings across all samples. The catalogue number and dilution range of the antibodies are listed in Table 2.

### Mitochondrial isolation

Mitochondria were isolated according to the method described earlier [78] and modified for the neuronal cultures. Briefly, DA neuron cell pellets were resuspended in mitochondria isolation buffer (MIB1: 225 mM Mannitol, 75 mM Sucrose, 5 mM HEPES, 1 mM EGTA and 1 mg/ml fatty acid free BSA) and homogenised using a ice-cold Dounce tissue grinder tube (appropriate for 1-1.5 ml homogenisation volume). The homogenate was centrifuged, and the mitochondrial pellet was washed and pellet down in MIB without BSA (MIB2) according to the protocol. The final pellet was resuspended in a very small volume of MIB2, and the protein concentration was determined using BCA Protein assay kit (Thermofisher Scientific, 23227) according to the manufacturer’s specifications.The isolated mitochondria were then used for respiratory measurements.

### Measurement of oxygen consumption rate

Measurements of mitochondrial respiration were conducted with the Seahorse Bioscience XFe96 bioanalyzer using the Seahorse XF Cell Mito Stress Test Kit (Agilent #103015-100). Final maturation of iPSCs into DA neurons were performed in XF96 cell culture microplates (Agilent #102416-100). On the day of the experiment, the culture medium was replaced with Seahorse XF Base medium (Agilent #103334-100) supplemented with 1 mM pyruvate (Gibco #11360070), 2 mM glutamine (Gibco #25030081) and 10 mM glucose (Gibco #A2494001) and incubated for 30 min at 37 °C in a CO_2_-free incubator before loading into the Seahorse Analyser. After measuring basal respiration, the drugs oligomycin (5 µM), FCCP (1.5 µM), and rotenone/antimycin A (0.5 µM/0.5 µM) were added to each well in sequential order. After the assay, cells were stained with Hoechst 33342 (5 µM; Thermo Scientific #62249) for 30 min. ImageXpress was then used to count the number of cell nuclei (cell numbers) in each well and normalised to get the basal respiration rate values.

The respiratory measurements of isolated mitochondria from mixed neuronal cultures were performed using a modified method described earlier[79] for Seahorse XFe96 cell culture plates assay. Briefly, 5 or 10 µg of mitochondrial protein was resuspended in 30 µL (1 well) of individual substrate mix (for example, pyruvate/malate substrate + mitochondria assay solution - MAS) and plated into each well. The cell culture plate was centrifuged at 2,000g for 20 minutes at 4 °C to form a uniform layer of mitochondria at the bottom. After centrifugation, 150 µL of substrate solution was carefully added to each well. Fresh injection solutions were made in MAS without BSA and loaded into the cartridge, and calibrated according to the manufacturer’s specifications. After calibration, the culture plate with mitochondria was inserted, and the assay was run essentially as described in the method.

### Transmission electron microscopy (TEM)

TEM was performed as previously described [76]. Briefly, fibroblasts and DA neurons grown on coverslips were fixed in electron microscopy (EM) fixative containing 2% glutaraldehyde (EMS, 16365) and 2% paraformaldehyde (EMS, 15710) in 0.1 M sodium cacodylate for 1 hour. Following fixation, cells were washed with 0.1 M cacodylate buffer (EMS, 11650) and then fixed in a solution of 1% osmium tetroxide (EMS, 19150) and 1% potassium ferricyanide (EMS, 25120-20) in 0.1 M sodium cacodylate. This was followed by sequential dehydration using ethanol. Coverslips were then embedded in epoxy resin (Araldite Kit, Agar Scientific Ltd., CY212) according to standard protocols. The embedded samples were sectioned into 50 nm slices using an ultramicrotome equipped with a diamond knife and mounted onto copper TEM grids. The grids were stained with lead citrate for 3 minutes before imaging. Images were captured using a Jeol 1400 Transmission Electron Microscope at magnifications ranging from 800× to 1200× (digital magnification). Images were imported into ImageJ/Fiji to quantify mitochondrial area, mitochondrial cristae density and mitochondrial aspect ratio. The autophagic vesicles were classified as autophagosomes, autolysosomes and lysosomes as per [24]. Briefly, in Fig 3D and Fig S3C, autophagosomes typically display clear double membranes or a single membrane, darker than the surrounding tissue in TEM images, with more circular shapes inside, indicating cargo (red arrows). Lysosomal membranes show highly organised inner folds, while lysosomal enzymes are characterised by a darker, more uniform appearance (green arrows). Autolysosomes are intermediate structures larger than lysosomes, frequently containing intact autophagosome contents (yellow arrows). Distinct dark spots (black arrows) may indicate protein aggregates; further work is needed to confirm their identity. Mitochondria and autophagic vesicles were manually traced to measure their areas, and the structures were verified by an Electron Microscopy expert. Quantification was performed in a double-blind manner.

### Quantification and statistical analysis

All statistical analyses were performed using GraphPad Prism (version10; GraphPad Software) and verified by a qualified statistician. Sample sizes (n) for each experiment represent values from at least 3 independent experiments.

For comparisons between two datasets, paired t-test or two-tailed unpaired t-tests were used for normally distributed data; when normality was not met, the Mann–Whitney test was used. For multi-group comparisons, one-way ANOVA was performed, followed by the Kruskal–Wallis test, with Dunn’s multiple comparisons correction applied. For experiments involving independent comparisons within the group, ordinary two-way ANOVA with Tukey’s multiple comparisons test (single pooled variance), Šidák multiple comparisons or Uncorrected Fisher’s LSD (single pooled variance) was used. Samples normalised to a reference value of 1 were analysed using one-sample Wilcoxon rank t-tests (Supplementary table 2).

Where multiple comparisons were conducted, adjusted p-values are reported following the appropriate post hoc correction. Adjusted p-values are provided in the graphs. Statistical significance was defined as adjusted p < 0.05.

The mitochondrial morphology parameters were heavily skewed, hence the values were log(Y) transformed before calculating the effect size, confidence intervals and unpaired t-test. Autophagic vesicle counts in neurons contained zeros; therefore, values were log(1+Y) transformed prior to two-way ANOVA. Effect sizes are reported as partial eta-squared (partial η²) with 95% confidence intervals.

Data are presented in the figures as mean ± standard deviation (SD). Effect sizes (for Mann-Whitney’s test comparison - Hedges’ g; Unpaired t-test – Cohen’s d; one-sample t-test - Glass’s Δ; ω² - for one-way ANOVA; partial η² - for two-way ANOVA). The noncentral F distribution method (for ANOVA) or the Bootstrap (all others) was used to compute 95% confidence intervals to aid interpretation of the magnitude and precision of observed effects. Wherever possible, individual data points and raw data distributions are shown in figures, and all raw numerical values used to generate graphs are provided in Supplementary Table 1. The statistical test, effect size, and confidence interval values for each graph are included in Supplementary Table 2 to promote transparency and reproducibility.

## Abbreviations

GBA1: Glucocerebrocidase
PD: Parkinson’s Disease
GCase: β-glucocerebrosidase enzyme
pH: potential of Hydrogen
V-ATPase: Vascular ATPase
iPSCs: induced Pluripotent Stem Cells
DA neuron: Dopaminergic neurons
OXPHOS: Oxidative Phosphorylation
ΔΨm: Mitochondrial Membrane Potential
TEM: Transmission Electron Microscopy
TMRM: TetraMethyl-Rhodamine Methyl ester
MTORC1: Mammalian Target of Rapamycin Complex I
FLIM-FRET: Fluorescence Lifetime Imaging (FLIM) with Förster Resonance Energy Transfer
NPs: Nanoparticles

## Declarations

## Ethics approval

The source and ethical approval committee for the iPSCs used in the study is tabulated in table 1.

## Consent for publication

Not applicable

## Availability of data and materials

All data generated or analysed during this study are included in this published article and its supplementary information files.

## Authors’ contributions

P.S., M.R.D - Conceptualisation, funding acquisition, investigation, visualization, methodology, writing–original draft, project administration, writing–review and editing. A.C.B –methodology, investigation, visualisation and data analysis. S.K, A.F, I.K, K.S and T.S.B– Methodology, investigation and data analysis. O.S, J.Z and M.G – Resources.

### Acknowledgements

The authors would like to thank the Michael J Fox Foundation and Parkinson’s UK Foundation for generously funding our project. The authors would also like to express their gratitude to the members of the Michael Duchen and Gyorgy Szabadkai labs for their feedback, as well as to undergraduate students, including Shail Bhatt and Oriane Marguet, for their assistance with the preliminary data. We acknowledge the MRC Centre for Neuromuscular Diseases Biobank (supported by the National Institute for Health Research Biomedical Research Centres at Great Ormond Street Hospital for Children, NHS Foundation Trust) for providing the age- and sex-matched healthy controls and GBA1-E326K fibroblasts used in this study. Figure 6 was created using Biorender.com. We thank Dr. Elizabeth Slavik-Smith from the Electron Microscopy facility at UCL, Division of Biosciences, for her assistance with EM. We are also grateful for the help provided by Dr. Christopher Von Ruhland, Facility Lead (Electron and Light Microscopy), Central Biotechnology Services, Cardiff University, with the identification of the autophagic vesicles. The authors thank Professor Peter Holmans, Division of Psychological Medicine and Clinical Neurosciences, Cardiff University, for his help and guidance with the statistical analysis.

